# Tracing cortical circuits in humans and non-human primates from high resolution connectomic, transcriptomic, and temporal dimensions

**DOI:** 10.1101/2021.04.30.442016

**Authors:** Christine J. Charvet, Kwadwo Ofori, Christine Baucum, Jianli Sun, Melinda S. Modrell, Khan Hekmatyar, Brian L. Edlow, Andre J. van der Kouwe

## Abstract

The neural circuits that support human cognition are a topic of enduring interest. Yet, the lack of tools available to map human brain circuits has precluded our ability to trace the human and non-human primate connectome. We harnessed high-resolution connectomic, anatomic, and transcriptomic data to investigate the evolution and development of frontal cortex circuitry. We applied machine learning to RNA sequencing data to find corresponding ages between humans and macaques and to compare the development of circuits across species. We transcriptionally defined neural circuits by testing for associations between gene expression and white matter maturation. We then considered transcriptional and structural growth to test whether frontal cortex circuit maturation is unusually extended in humans relative to other species. We also considered gene expression and high-resolution diffusion MR tractography of adult brains to test for cross-species variation in frontal cortex circuits. We found that frontal cortex circuitry development is extended in primates, and concomitant with an expansion in cortico-cortical pathways compared with mice in adulthood. Importantly, we found that these parameters varied relatively little across humans and studied primates. These data identify a surprising collection of conserved features in frontal cortex circuits across humans and Old World monkeys. Our work demonstrates that integrating transcriptional and connectomic data across temporal dimensions is a robust approach to trace the evolution of brain connectomics in primates.

**Significance Statement:** We lack appropriate tools to visualize the human brain connectome. We develop new approaches to study connections in the human and non-human primate brains. The integration of transcription with structure offers an unprecedented opportunity to study circuitry evolution. Our integrative approach finds corresponding ages across species and transcriptionally defines neural circuits. We used this information to test for variation in circuit maturation across species and found a surprising constellation of similar features in frontal cortex neural circuits across humans and primates. Integrating across scales of biological organization expands the repertoire of tools available to study connections in primates, which opens new avenues to study connections in health and diseases of the human brain.

## Introduction

The human frontal cortex (FC) is larger than that of many other species. The claim that the FC, and particularly the prefrontal cortex (PFC) white matter, is unusually large in humans compared with other primates is not without controversy (1-3). The expansion of the PFC white matter points to possible modifications in neural circuits across species. However, the lack of available tools has precluded our ability to visualize connections in the human brain (4). At the microscale, RNA sequencing from bulk samples and single cells offers an unprecedented perspective on the classification of cell types and transcriptional profiles. These data, however, lack key information about the structure of circuits (5-9). At the macroscale, diffusion MR tractography offers an exquisite three-dimensional perspective of pathways across the brain (10-15), which makes diffusion MR tractography an exciting tool to explore the connectome. The human brain scans we used in this study are of unprecedentedly high resolution for the study of primate brain connectomics (14, 15), yet the termination sites of tracts can be imprecise. Consequently, results from tractography necessitate validation (16, 17). We harnessed transcriptomic and connectomic data across temporal scales to trace the evolution of connections in the primate brain (18-20). We show that the integration of RNA sequencing with diffusion MR tractography is a robust approach to study connections.

Neuronal populations across the depth of the cortex can be used to predict connectivity patterns (21-22). For example, the number of excitatory layer III FC neurons, which preferentially form long-range cross-cortical projections, is increased in humans relative to mice (5, 23-24). These observations suggest a major expansion in long-range cortico-cortical pathways in humans relative to mice. Comparative analyses of diffusion MR tractography have likewise revealed an expansion of cortico-cortical fibers in primates relative to rodents (23). Several genes, such as some supragranular-enriched (SE) genes, identify long-range projecting (LRP) neurons (e.g., *NEFH, VAMP1, SCN4B*) in layers II-III and V-VI. However, a major hurdle in linking transcription with tractography is the lack of adequate markers to transcriptionally define neurons that project over long distances (18, 24). To overcome this hurdle, we aligned transcriptional and structural variation over the course of human development to transcriptionally define neurons that project through the white matter. We focused our attention on the evolution of cortico-cortical pathways given major reported differences between humans and mice (5, 23-24).

Developmental timelines vary tremendously across species, with human development extended compared with many other model systems (4, 25, 26). Because we lack appropriate norming procedures to compare brain development across species, we applied machine learning to RNA sequencing data to align ages in humans and macaques. We then tested whether FC maturation is unusually extended in humans relative to macaques after accounting for variation in developmental schedules (25, 26, 27). These data reveal that the transcriptional and structural features of FC circuits, presumed to be unique to humans, are in fact shared with macaques.

## Results

We used structural and transcriptional variation to find corresponding ages across species. We then integrated these data to test for cross-species variation in FC neural circuits.

### Corresponding Time Points during Postnatal Development

We used time points to find corresponding ages between humans and macaques (n=96; Table S1). These data included time points from a glmnet model applied to normalized gene expression in reads per kilobase per million (RPKM) from human and macaque PFC of different ages (n=38 humans, n=31 macaques; Fig. 1 and *SI Appendix*, Fig. S1; (28)). We selected genes with minimum expression averaged across samples (i.e., log10(RPKM)>1; n=8,014). We did not predict ages beyond 10.6 years in macaques because samples at late stages are sparse (*SI Appendix*, Fig. S1). We trained a glmnet model to predict ages from normalized gene expression in humans (cross-validation=10; n=38). This model has high predictive accuracy because sampling 70% of the data to train the model (R^2^=0.99) resulted in strong correlations between log-10 transformed predicted and observed ages (R^2^=0.98).

**Figure 1.**
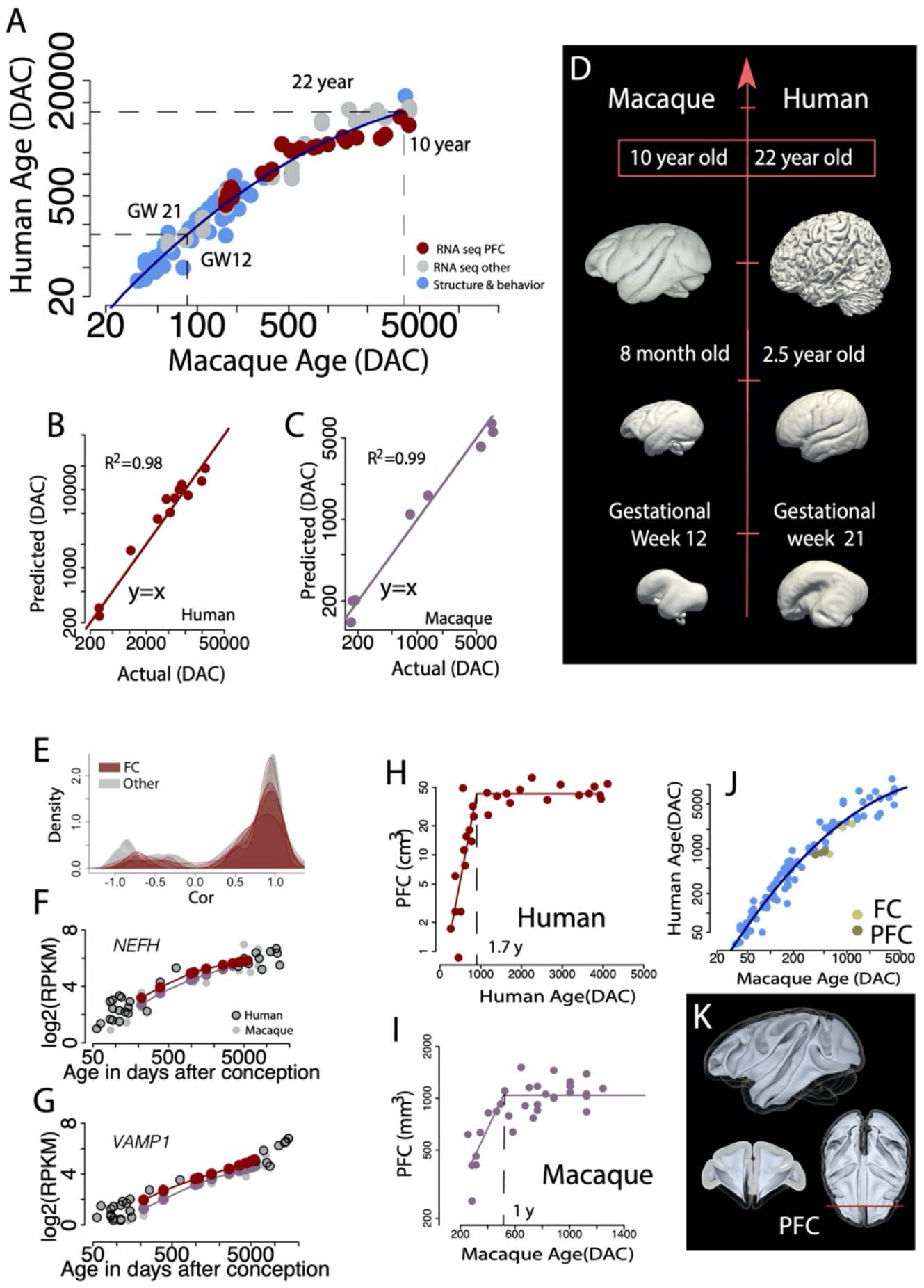
(*A*) We identified corresponding ages between humans and macaques from abrupt and gradual changes in structural and transcriptional variation. We extracted corresponding time points by training a glmnet model to predict age from RNA sequencing data in humans and subsequently predicted ages from normalized gene expression in macaques. Ages are expressed in days after conception (DAC), unless otherwise noted. (*B* and *C*) Models trained to predict age in each species have a high accuracy as evidenced by the strong correlation between predicted and observed values in humans (*B*) as in macaques (*C*). (*D*) With these data, we identified corresponding ages across fetal and postnatal time points between humans and macaques. For example, a macaque at gestational week (GW) 12 is equivalent to a human at GW 21, and a 10-year-old macaque is equivalent to a 22-year-old human. We then addressed whether the development of FC circuits is protracted in humans relative to macaques after controlling for variation in developmental schedules. (*E–G*) We tested whether temporal trajectories in SE gene expression and (*H*–*K*) growth trajectories in the FC and PFC white matter are extended in humans relative to macaques after aligning ages. We mapped age in macaques to that of humans and we tested whether the temporal trajectories of SE genes in the FC are significantly different relative to other cortical areas in humans and macaque. (*E*) Correlation coefficients from the FC are not significantly different from those from other cortical areas. (*F* and *G*) SE gene expression of *NEFH* and *VAMP1* in the dorsolateral frontal cortex of humans and macaques overlap extensively in the two species. (*H–J*) We extracted epochs from growth trajectories in the PFC and FC white matter in humans and macaques. (*H* and *I*) We then fit non-linear regressions from the growth trajectories, as shown here for the PFC white matter, which shows it largely ceases to grow at 1.7 years of age in humans and 1 year of age in macaques. (*J*) Time points extracted from these growth trajectories overlap with other time points. (*K*) The PFC white matter was defined as the white matter anterior to the corpus callosum as highlighted on a macaque template. The red vertical line illustrates the posterior boundary of the PFC white matter. These data indicate that FC growth is not protracted in humans relative to macaques. Smooth surfaces of MR scans are from (36, 42-47) and the Allen Brain Atlas. (Note: the ages in *D* do not correspond exactly to the labeled age).

The same approach applied to normalized gene expression in macaques predicted age of macaques (R^2^=0.99). We then used a glmnet model (R^2^=0.97; lambda=0.053) trained from human samples (n=38) to predict ages from normalized gene expression in macaques. This approach yielded 23 corresponding time points in humans and macaques. We fit a quadratic model on the log-transformed time points in humans and macaques (y= −1.96+ 2.74x-0.31x^2^; R^2^=0.95, n=96) to find corresponding ages across fetal and postnatal time points. Early in development, ages are roughly similar in the two species but time points occur much later in humans than in macaques by adulthood (Fig. 1). We next tested whether FC development at the transcriptional and the structural level (Figs. 1 and 2) is unusually extended in humans.

**Figure 2.**
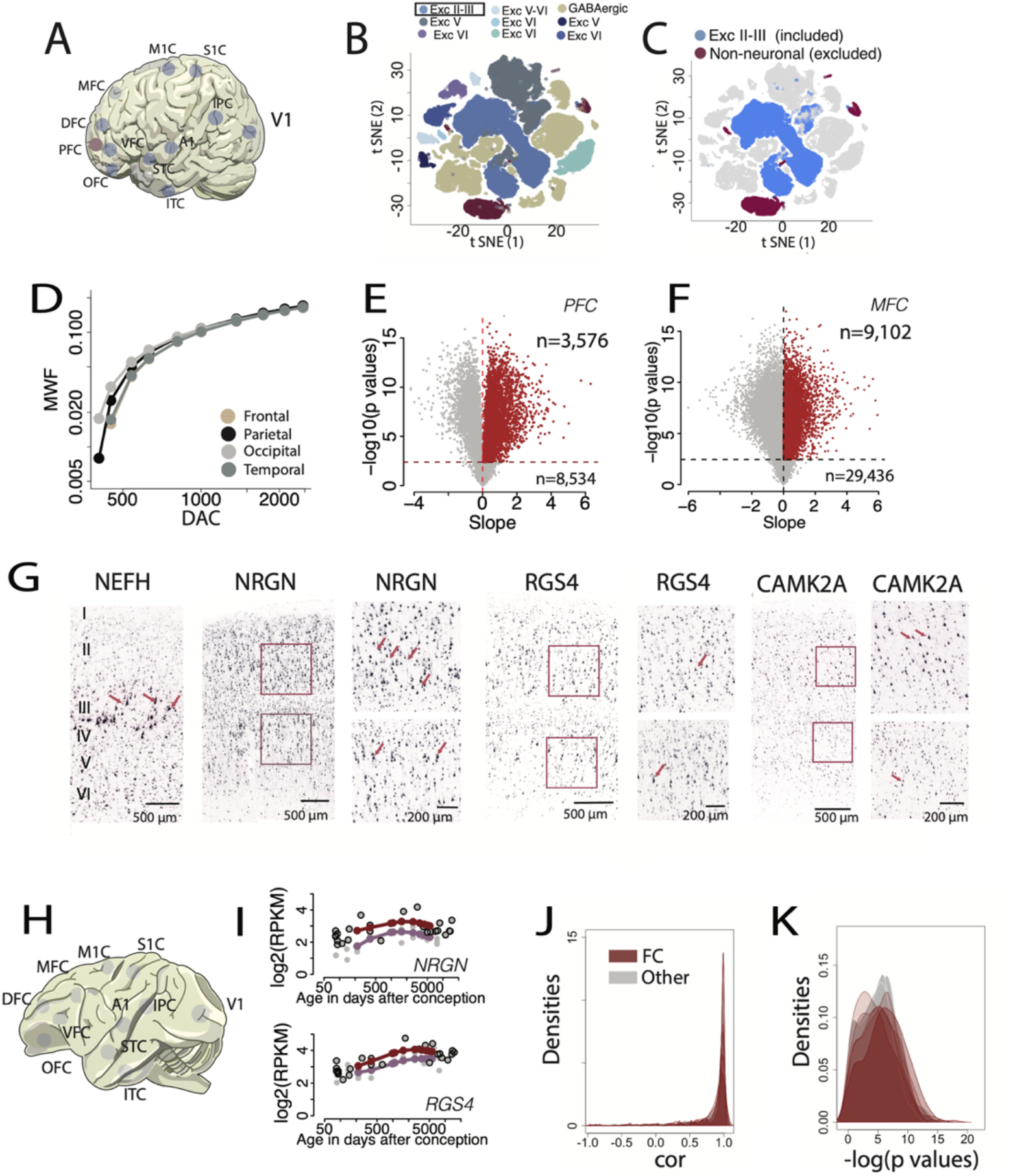
We identified 250 LRP markers by systematically testing for variation in transcription and myelination across cortical areas. (*A*) We considered gene expression across multiple areas and data-sets as highlighted on a schematic of a human brain (8, 28). (*B* and *C*) We used a single cell RNA sequencing dataset where clusters of cell populations were defined with tSNE distributed stochastic neighbor embedding (30) to filter candidate genes by cell type. (*C*) Our list of LRP markers includes genes that are expressed by layer II-III neurons but excludes those expressed by non-neuronal cells. (*D*) We extracted MWF values from equations of different lobes. (*E–G*) We fit a smooth spline through the log10(RPKM) values versus age expressed in days after conception (DAC) to extrapolate normalized gene expression, then we systematically tested associations between myelination across datasets. Two examples of association are shown in *E* (for the PFC) and *F* (for the MFC) using the data (28), and the Allen Brain atlas, respectively. We selected significant and positive (slope>0) associations after correcting for multiple testing (BY: p<0.05). (*G*) Expression of LRP markers (*NEFH, NRGN, RGS4*, and *CAMK2A*) in pyramidal neurons (arrows) indicate this approach is valid to identify LRP markers. Boxes indicate regions shown in higher power views. (*H–K*) We considered temporal variation in transcription across the cortex of humans and macaques (*H*) to test for variation in the development of FC circuitry relative to other cortical areas in both species. (*I*) We fit a smooth spline through normalized gene expression versus age in humans and macaques as exemplified in the primary motor cortex (M1). We then extracted normalized gene expression from smooth splines at corresponding ages across humans and macaques (n=10). Shown in this example are the results for the genes *NRGN* and *RGS4*. (We correlated the expression of LRP markers across humans and macaques for each cortical area and tested for differences in correlation coefficients (*J*) and evaluated significance tests (*K*). No significant differences in correlation coefficients are observed between the FC and other cortical areas. Overall, we find conservation in the developmental time course of LRP marker expression in the human FC relative to macaques. Additional abbreviations: A1: primary auditory cortex; DFC: dorsolateral frontal cortex; IPC: inferior parietal cortex; ITC: inferior temporal cortex; MFC: medial frontal cortex; M1C: primary motor cortex; OFC: orbitofrontal cortex; PFC: prefrontal cortex; S1C: primary somatosensory cortex STC: superior temporal cortex; VFC: ventral frontal cortex; V1: primary visual cortex.

### Transcriptional-defined Neural Circuits to Detect Variation in Neural Circuit Maturation

We used the expression of SE genes (n=16) and LRP markers to test for cross-species variation in FC circuit development (Figs. 1 and 2, *SI Appendix*, Figs. S2–S4; Table S2). We mapped age in macaques onto humans, extrapolated normalized gene expression at corresponding time points in both species, and then correlated SE gene expression in humans and macaques (Fig. 1). We found that these correlation coefficients are not significantly different across cortical areas (ANOVA: F=0.145; p=0.99; n=176). Although these findings suggest conservation in FC circuit maturation, it is not clear whether genes other than SE genes are best suited to track FC circuit maturation across species.

We identified LRP markers by testing for associations between white matter maturation and gene expression (Fig. 2; (8, 29) across cortical areas. Genes were filtered to have minimum expression averaged across regions and ages (log10(RPKM)>1). We extrapolated myelin water fraction (MWF) values from previous work (29) (Table S3). We fit a smooth spline through log-based 10(RPKM) values versus age (i.e., log-10 days after conception) to extrapolate gene expression and MWF at matching ages (n=10) and across cortical areas (n=10). We iteratively tested for associations between gene expression and MWF across cortical areas with a linear model to normalized gene expression. We selected significant and positive associations (slope>0) after correcting for multiple testing (Benjamini & Yekutieli: BY; p<05). We used single-cell RNA sequencing (30) from the motor cortex to exclude genes expressed by non-neuronal cells but include those expressed by layer II-III neurons (*SI Appendix*, Table S4). These filtering steps resulted in 250 LRP markers. These genes are expressed by large pyramidal neurons, which increase in expression postnatally, and a subset of these are SE genes (e.g., *NEFH, VAMP1*)(*SI Appendix*, Table S5 Figs. S2–S4). These observations support the validity of these genes as LRP markers

We tested whether temporal profiles in the expression of these LRP markers are different between humans and macaques. We fit a smooth spline through the log2(RPKM) values versus log-10 ages to extrapolate gene expression at equivalent ages in macaques and humans (n=10). We correlated the expression of LRP markers (n=234) over the course of development and across cortical areas (n=11). An ANOVA on the correlation coefficients pointed to significant differences across cortical areas (ANOVA: F=3.12, p<0.01; n=2,574), but post hoc Tukey HSD tests showed that correlation coefficients from the FC areas were not significantly different (significance threshold set to p<0.01) from those of other cortical areas (Fig. 2; statistics in *SI Appendix*, Table S6). These data showcase the strong similarity in FC development between humans and macaques.

### Structural Variation in FC Circuit Development

We compared white matter growth across species to test whether LRP neuron maturation is unusually extended in humans (Fig. 1 and *SI Appendix*, Fig. S6). We fit non-linear regressions with age in days after conception as the predictor variable and the FC and log10-based PFC white matter volume as the dependent variable (Fig. 1; see *SI Appendix*, Table S7; Figs. S5 and S6). We captured the ages at which the FC and PFC white matter reach percentages of adult volumes (e.g., 80%, 90%, 100%; *SI Appendix*, Fig. S5). We found that these time points overlap with other time points (Fig. 1). For example, the PFC white matter ceases to grow at 1.74 years of age in humans (R^2^=0.76, n=28, p<0.001) and at ∼1 year of age in macaques (R^2^=0.66, n=32, p<0.001). An ANOVA showed that the addition of time points from these growth trajectories did not account for a significant percentage (p=0.24) of the variance (F=1,355, p<0.05; n=102). We subsampled these data and showed that the age of growth cessation was largely invariant with respect to sample size, demonstrating that these results were not driven by outliers (*SI Appendix*, Fig. S6). We found no evidence for protracted FC white matter maturation in humans.

### Cross-species Variation in Tractography of Adult FC Circuits

We used diffusion MR tractography to test for modifications to FC connectivity across humans, macaques, and mice (Figs. 3 and 4; *SI Appendix*, Figs. S7–S14, Tables S8-S10). We developed a new approach to quantify pathways. In developing this approach, we compared the orientation and location of fibers from tractography, tract-tracers, immunohistochemical markers, and myelin stains to assess sites of potential inaccuracies in the tractography (Fig. 4 and *SI Appendix*, Figs. S7– S14). In mice, the direction and location of tracts within the white matter were concordant across methods but tracking accuracy appeared limited at the grey to white matter boundary. We therefore classified pathways based on the orientation and direction of pathways within the white matter (Fig. 4). We placed voxels in spaced sections through the FC white matter and we classified pathways according to their orientation (Figs. 3 and 4). The percentage of cortico-cortical pathways was significantly greater in the primate FC relative to mice but no significant differences were observed between humans and macaques (ANOVA: F=14.82, p<0.01; n=14). Overall, we found no significant differences in the relative proportion of FC and PFC pathway types across primates (t tests; Fig. 3, statistics in *SI Appendix*, Table S8). Subcortical pathways were significantly expanded in the PFC of humans versus macaques (t=5.1, p<0.008). Relative FC pathway types were relatively invariant with respect to sampling, tested directions, pathway reconstruction, and resolution (*SI Appendix*, Figs. S12–S14). Although the cross-species variation in pathway types or lack thereof were robust to variations in imaging parameters, the limitations inherent to tractography led us to generate evidence from multiple scales to ensure the accuracy of these findings.

**Figure 3.**
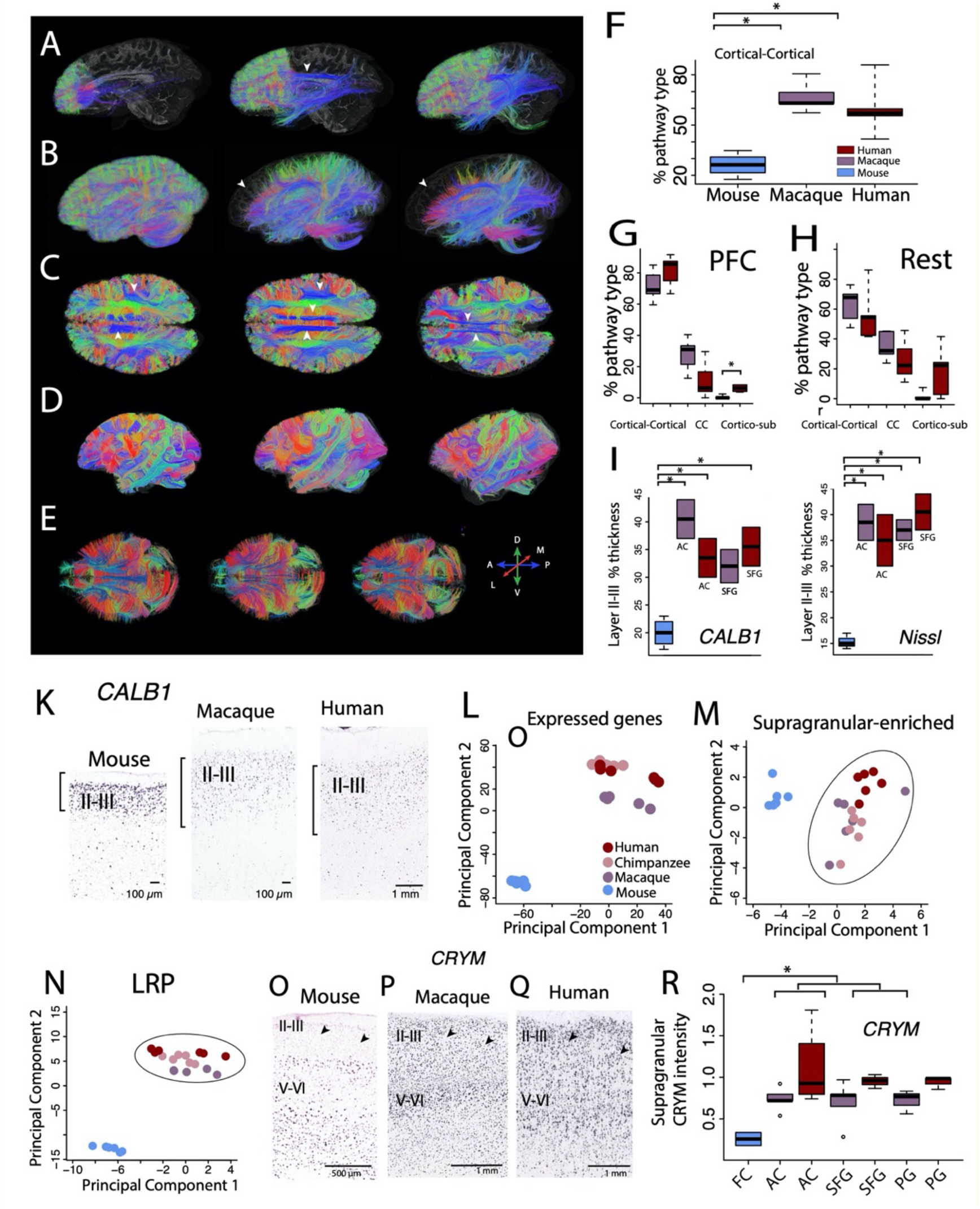
Analysis of pathways with tractography, cytoarchitecture, and transcription reveal modifications in adult FC circuits across humans, macaques, and mice. (*A*) The diffusion MR scans, which are of high resolution, identify pathways coursing through the human brain. (*B*) Brain tractography in humans with variable minimum length thresholds (0, 20, and 40 mm) reveal different pathways across the cortex. In particular, there appears to be preponderance of short fibers within the FC as evidenced by a lack of tracts within the anterior cortex (white arrowheads) when 20 or 40 mm minimum length thresholds are set. (*C*) Horizontal slices (1-mm wide) show pathways coursing across the white matter, some of which span the FC (e.g., cingulate cortex, arcuate fasciculus; arrowheads). Diffusion MR scans of (*D*) primate and (*E*) mouse brains were also used to classify pathway types. (*F*–*H*) Percentage of pathway types in the FC white matter of humans, macaques, and mice show that the relative percentage of cortico-cortical pathways is significantly greater in humans and macaques compared with mice. (*: p < 0.05). No significant differences in pathway type are detected between humans and macaques whether we consider the PFC (*G*), or the rest of the FC (*H*). (*I–R*) The relative thickness of layer II-III as defined by *CALB1* expression (*I* and *K*) and Nissl stains (*J*) is significantly greater in the primate FC compared with mice but not between humans and macaques. This is true whether we consider the anterior cingulate cortex (AC) or the superior frontal gyrus (SFG). (*L*–*N*) PCAs testing whether transcriptional profiles of LRP or SE genes differ between humans and macaques. A PCA of orthologous expressed genes (*L*) shows that samples cluster by species but PCAs of expressed SE genes (*M*) and LRP markers (*N*) show that macaque samples cluster with those of humans and chimpanzees. These observations support the notion that transcriptional profiles of LRP neurons are highly similar between humans and macaques. (*O–Q*) *CRYM* (a SE gene) expression is higher in supragranular layers of humans and macaque superior frontal gyrus (SFG) compared with mice (arrowheads). (*R*) The expression of *CRYM* is significantly higher in primate supragranular layers compared with the FC of mice, but no significant differences were observed between humans and macaques. This is true whether we compare the mouse FC with the anterior cingulate cortex (AC), the superior frontal gyrus (SFG), or the precentral gyrus (PG) of macaques and humans. Humans and macaques share a constellation of conserved features in FC neural circuits.

**Figure 4.**
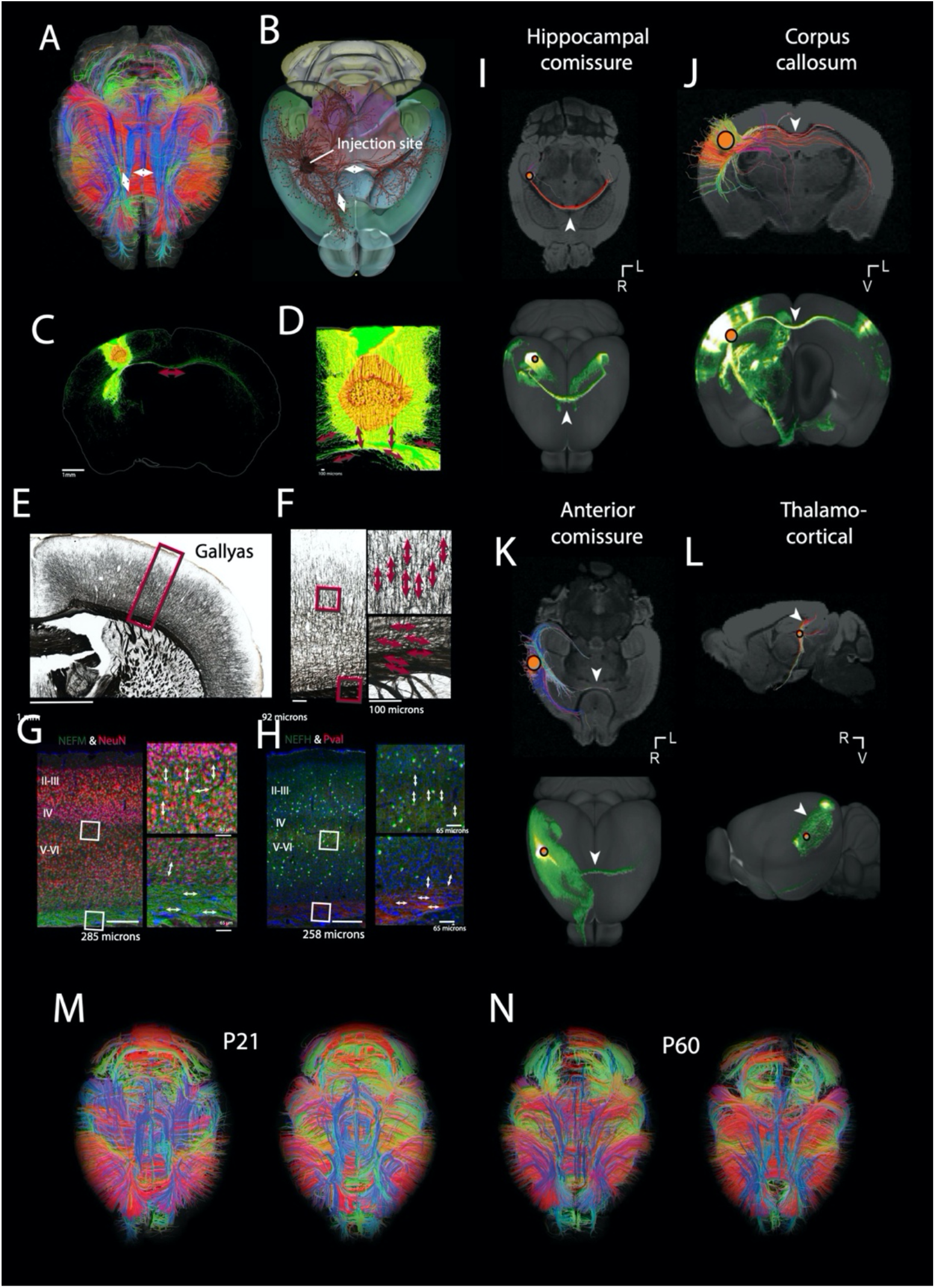
We compared tractography with tract-tracers and histological stains to develop approaches to quantify pathway types. In mice, white matter pathways penetrate the grey matter at roughly right angles. We refrained from using endpoints within the grey matter as a basis for classification because of the potentially reduced accuracy of the tractography at the grey to white boundary. We compared diffusion MR tractography (*A*) with tract-tracers (*B–D*), myelin (*E* and *F*), and immunohistochemical stains (*G* and *H*) to visualize the orientation of small to large axons within the grey and white matter. Tract-tracers injected in the primary somatosensory area (upper limb-D) show fibers radially aligned within the grey matter. At the intersection of the grey and white matter, these fibers make a sharp turn and course across the medial to lateral axes, towards subcortical structures, and rostrally (white arrows; *A* and *B*). (*E* and *F*) Gallyas stains show myelinated axons course radially within the grey matter but course preferentially along the medial to the lateral axis in the white matter (red boxes show close-up views). Antibodies against neurofilament medium polypeptide (*NEFM*) and neurofilament heavy polypeptide (NEFH), which label neurons of small (*G*) and large calibers (*H*), respectively, further support the observations from diffusion MR tractography and the gallyas stain. That is, NEFH+ and NEFM+ neurons are oriented radially within the grey matter but course across the medial to lateral axis in the white matter; white boxes highlight the close up views shown in *G* and *H*. Neun and Pval label neurons. These observations suggest that large and small caliber axons make a sharp turn at the grey to white matter boundary. Sharp turns at the grey to white matter boundary are associated with reduced accuracy. Fiber orientation determined from tract-tracers and tractography are concordant across methods (statistics in *SI Appendix*, Table S14). (*I–L*) ROIs were used to capture well-known pathways coursing through the white matter (*I*: hippocampal commissure, *J*: corpus callosum, *K*: anterior commissure, *L*: thalamocortical fibers from the lateral geniculate nucleus to the primary visual cortex, white arrow-heads). Tract-tracer data are from the Allen Brain Institute Mouse Connectivity Database. (*M* and *N*) Examples of diffusion MR scans in mice reveal consistency in fiber tracking across brains at P60 and P21.

### Cross-species Variation in Transcription of Adult FC Circuits

Given that FC cortico-cortical pathways are expanded in primates, we focused on layer II-III neurons, many of which form cortico-cortical projections. First, we considered the cytoarchitecture of the layer to test for modifications in FC long-range projecting fibers across mice, macaques, and humans (Fig. 3). We observed that layer II-III in the FC, as defined transcriptionally (i.e., *CALB1* expression) and cyto-architecturally, is relatively expanded in humans and macaques compared with mice. This was true whether we considered the anterior cingulate (*CALB1*: ANOVA: F=22.72; p<0.01, n=8; Nissl: F=20.17; p<0.01) or the superior frontal gyrus (*CALB1*: ANOVA: F=15.62; p<0.01, n=8; Nissl: F=15.16; p<0.01; Fig. 3I). Post-hoc Tukey HSD tests showed the relative thickness of FC layer II-III is significantly different between primates and mice but not between humans and macaques (Fig. 3; statistics in *SI Appendix*, Table S12 and 13). Moreover, the expansion of layer II-III in macaques is concomitant with an expansion of SE gene expression in the layer (*SI Appendix*, Figs. S15-–S18). Our results showed that there are major differences in layer II-III between primates and mice but strong similarity across humans and macaques.

We next considered transcriptional variation of SE and LRP markers to investigate cross-species variation in adult FC circuits (Fig. 3 and *SI Appendix*, S15–S18). In a principal component analysis (PCA) applied to log-transformed expressed orthologous genes, genes clustered according to species (Fig. 3L; first 3 principal components (PCs): 71.77% of the variance; n=10,682). In PCAs on the log-transformed expressed SE genes (Fig. 3M; first 3 PCs: 73.9% of the variance; n=17) and LRP markers (Fig. 3N; first 3 PCs: 77.07%; n=235), primates clustered together, demonstrating the strong similarity in expression of SE and LRP genes between humans and macaques. We also evaluated *CRYM* expression (i.e., an SE gene) to confirm similarities in SE gene expression across humans and macaques (Fig. 3O–R and *SI Appendix*, Fig. S17, Table S12). We quantified the relative expression of *CRYM* across supra- and infragranular layers in humans, macaques, and mice and found that expression patterns varied across species whether we compared the mouse FC with the superior frontal gyrus (ANOVA: F=8.94; n=11; p<0.01), the anterior cingulate (F=4.86; n=11; p<0.01), or the precentral gyrus (F=22.04; n=8; p<0.01) of macaques and humans (Fig. 3R). Tukey HSD tests showed no significant differences in the expression profile of *CRYM* between humans and macaques but differences between primates and mice (*SI Appendix*, Table S13). These data showcase the strong similarities in FC neural circuitry between humans and macaques.

## Discussion

The integration of transcription with neuroimaging is an effective approach to identify conservation and variation in biological programs linked to modifications of circuits in human evolution. This integrative approach identified corresponding ages, and tested for FC neural circuit variation across species, which creates novel opportunities to study circuits in human health and disease, and across species.

### Corresponding Ages from Transcriptional and Structural Variation

We identified corresponding postnatal ages in humans and macaques. This work builds on a previous line of work called the Translating Time Project (www.translatingtime.org), which relied on abrupt changes that unfold during development to find corresponding ages across model organisms and humans (8, 25-27). We collected 354 transformations across 19 mammalian model organisms to find corresponding time points during prenatal development. Extracting time points from gradual changes in transcription and structure, in addition to abrupt transformations, reveals corresponding postnatal ages (8, 23). Each metric may have uncertainties, but the use of multiple metrics ensures the robust determination of corresponding ages across species.

### Limitations and Opportunities for Diffusion MR Imaging

The integration of transcription with diffusion MR tractography permits tracing the evolution of FC neural circuits. Diffusion MR tractography reveals a three-dimensional perspective of pathways (12-14), However, the sole use of tractography to trace neural circuits is problematic because of its limited ability to resolve crossing fibers or locate tract termination sites (Fig. 4 I-J, K-L). Considering these caveats, we compared FC pathway types coursing through the white matter. We selected this approach because of our comparisons of tractography with tract-tracers, immunohistochemistry, and myelin stains. The orientation and location of fibers aligned with observations from histology and tract-tracers (Fig. 4). Our analyses, which were designed to overcome limitations of diffusion MR tractography, withstood variations in sampling, direction, and resolution (Figs. S13 and S14). Nevertheless, we still lack methods to ensure the accuracy of diffusion MR tractography (10, 31). The lack of alternative methods to map connections motivated us to integrate transcriptomic and structural information to rigorously trace connections across species.

### Enhanced Methods to study FC Circuits in Primates Reveal Conservation

Although RNA sequencing from bulk and single cells offers an unprecedented perspective to track developmental programs, these metrics lack key information about the structural composition of circuits. There is often a lack of one-to-one correspondence between projection patterns and gene expression (32). We, therefore, identified LRP markers by aligning structural and transcriptomic variation during human development. This novel approach was instrumental in systematically identifying genes expressed by LRP neurons. The concordance of findings from transcription and neuroimaging enabled tracing modifications to circuits in primates.

We drew from multiple lines of evidence to test for modifications in FC circuitry across species. We compared trajectories in transcriptional profiles of LRP neurons, white matter maturation, pathway types, and transcription across layers. Testing for differences across these scales revealed no evidence that human FC circuits are unusual relative to macaques. Rather, these results systematically highlighted conservation in FC circuits across humans and other primates (8, 33). Past work considered white matter volumes or transcriptional information in isolation to assess whether FC circuits are unusual in humans but these studies did not reach a consensus. We show that tracing human FC circuits from connectomic, transcriptomic, and temporal dimensions moves us forward in mapping circuits in the human and non-human primate brain.

## Materials and Methods

We discuss how we found corresponding ages across species and how we tested for cross-species modifications in FC circuitry. All statistics were performed with the programming language R. All ages were expressed in days after conception.

### Transcriptional and Structural Variation to Infer Corresponding Ages Across Species

We used 104 time points to find corresponding ages between humans and macaques (Fig. 1 and *SI Appendix*, Table S1, Fig. S1; 28). These included time points extracted from an RNA sequencing dataset from the PFC of humans and macaques. We selected genes with log-based 10 expressions in RPKM>1 averaged across samples per species. We fit a glmnet model (cross-validation: n=10; repeat=5, tune length=5) to log10(RPKM**+1**) in humans. We then used this model to predict ages from normalized gene expression in macaques. We considered this model accurate because ages predicted from these and other time points accounted for 95% of the variance when nonlinear regressions were fit to these data (Fig. 1). Due to possible variation in extrapolating ages from machine learning tools, the time points from the glmnet model were included with other time points to find corresponding ages across species.

### Transcriptional Definition of Cortical Long-Range Projecting Neurons

We used gene expression data to test for modifications to FC circuits. We considered SE genes because of differences in expression patterns between humans and mice, and because they are expressed in layers III–V where neuronal somas that project over long distances are located (34). It is, however, not clear whether other genes are better suited to study LRP neurons. We therefore developed novel approaches to systematically identify LRP markers by aligning temporal variation in gene expression and structure.

We identified LRP markers by testing for associations between transcription and maturation over the course of human development. We considered MWF as an index of white matter maturation across lobes (29) (*SI Appendix*, Table S3) and used multiple RNA sequencing datasets (8, 28) (Fig. S1). LRP markers were defined as genes 1) that were expressed by layer II-III neurons but not non-neuronal cells, and 2) that have expression patterns significantly associated with MWF. We added a value of 1 before logging the expression of each sample to consider genes that may not be expressed at a specific age. We fit smooth splines (degrees of freedom=4) through log10(RPKM) values versus age to extrapolate data at matching ages (n=10; from 405 DAC to 6 years of age). Only expression profiles that significantly associated with MWF across all tested areas were considered LRP markers. We used single-nucleus transcriptomes from the human primary motor cortex (n=2; 18-68 years, n=76,535 nuclei) (30) to filter these genes by cell types (Table S4). An expressed gene was defined by a count>0. We also used in situ hybridization from the Allen Brain Atlas dataset to evaluate the spatial expression of LRP markers and SE genes in order to test whether FC neural circuitry maturation is unusual in humans relative to macaques. We considered RNA sequencing data from macaques (n=26) and humans (n=36) and across cortical areas (n=11) (8). We translated age in macaques to that of humans and fit smooth splines (degrees of freedom=4) to compare normalized gene expression across these two species.

### Structural MR Scans to Test for Variation in FC White Matter Maturation

The white matter houses long-range projections. We compared white matter growth to test for variation in the timeline of FC circuits across species. Our definitions of FC and PFC follow those used previously (19, 26). Structural MR scans of macaque brains were obtained from the UNC-Wisconsin Rhesus Macaque Neurodevelopment database (n=32; *SI Appendix*, Table S11). We used Fiji to measure the PFC white matter, which was defined as white matter anterior to the corpus callosum, consistent with previous definitions (Fig. 1 and *SI Appendix*, S5). For this, we used data generated for this study other studies (26, 33, 35). We measured the PFC white matter area across sections (in at least every other section) using an approach similar to that used previously (26). We reconstructed volumes by multiplying the area, section thickness, and section spacing. Nonlinear regressions (library easynls, model=3) were used to detect the age of growth cessation (Fig. 2 and *SI Appendix*, Figs. S5 and S6) with the caveat that some growth may persist beyond identified time points.

### Diffusion MR Imaging Protocols and Tractography

We used diffusion MRI datasets of 14 individuals: humans n=5), mice (n=4), macaques (n=4), and Sykes monkey(n=1). Some of these datasets were previously collected (e.g., Japanese Monkey Brain Center; 36-38). We used diffusion MR datasets of human brains (n=4) scanned on a 3T Siemens Tim Trio scanner with a 32-channel head coil at the Massachusetts General Hospital Athinoula A. Martinos Center for Biomedical Imaging. The resolution of the human MR scans was 0.75 mm isotropic (diffusion MRI data acquisition duration=∼31 hours). Diffusion-weighted data were acquired with a 3D steady-state free precession sequence. Diffusion weighting was applied along 90 directions distributed over the unit sphere (effective b value = 4,080 s/mm2; 12 b0s; TR∼28.87ms, TE∼24.44ms). We collected diffusion MR scans of mouse brains (n=8) at postnatal day (P) 21 and P 60 using a 9.4T Bruker scanner at the University of Delaware (Fig. 4 and *SI Appendix*, Figs. S9 and S14). A three-dimensional diffusion-weighted spin-echo echo planar imaging (SE-EPI) sequence (TR of ∼500 ms, TE∼40 ms; resolution: 100 um isotropic) was used to image the mouse brains. Sixty diffusion-weighted measurements (b=4,000 s/mm2) and non-diffusion-weighted measurements (5 b=0s) were acquired.

### Tractography Quantification and Comparison with Tract-tracers

The accuracy of diffusion MR tractography has remained elusive due to a lack of alternative tools to map the connectome in humans. We compared diffusion MR tractography with EGFP injections to assess which metrics should be extracted from tractography in mice (Fig. 4 and *SI Appendix*, Fig. S13). Given that the accuracy of tractography appeared compromised at the grey-white matter boundary (Fig. 4 and *SI Appendix*, Fig. S11), we randomly selected voxels across spaced planes along the anterior to posterior axis of the FC white matter, with planes varying across individuals. We then classified pathways based on orientation and direction within the white matter. Fibers were classified as those belonging to the corpus callosum, cingulate bundle, other cortico-cortical, or subcortical-cortical pathways (*SI Appendix*, Fig. S14 and Table S10). If a pathway was observed coursing through the dorsal midline, it was considered callosal regardless of its terminations. Pathways connecting the cortical and lateral limbic structures were considered cortico-subcortical pathways. U fibers or long-range pathways connecting cortical areas (e.g., arcuate fasciculus) were considered cortico-cortical. We focused on cortico-cortical pathways because of the expansion of layer II-III in primates relative to rodents. We tested how sampling, resolution, and imaging protocols impacted pathway types (*SI Appendix*, Figs. S12–14). We observed that high angular resolution diffusion imaging (HARDI) and diffusion tensor imaging (DTI) reconstruction yielded comparable results in the mouse FC at P60 (y=0.82x+4.52; R^2^=0.89; p<0.01). We also tested for temporal variation in the relative percentage of pathway types of mouse brains at P21 versus P60. An ANOVA with age and pathway types as factors showed trends but no signific effect for age (p<0.05) on the pathway types (F=13.16, p<0.01; n=32). Frontal cortex white matter growth in mice ceases before P60 (19); therefore, P60 should represent adult proportions in pathway types.

### Comparative Analyses of Supragranular-Enriched Gene expression in the FC

We leveraged RNA sequencing datasets from bulk samples (n=24) (39-42), single cells, and in situ hybridization (ISH) to detect cross-species variation in SE and LRP gene expression across the human, macaque, and mouse FC (*SI Appendix*, Table S2). We performed PCAs on these data, and measured gene expression intensity across layers as we had done previously (23, 26) (*SI Appendix*).

## Data Availability

Data and scripts will be available on dryad.

## Acknowledgments

We thank Rohina Niamat and Drs. Harrington, Whitaker, and Halley for their help. Images were taken from the Allen Institute website and the Brainspan Atlas of the developing human brain. Data are available at http://www.brainspan.org and http://developingmouse.brain-map.org, which is supported by the NIH Contract HHSN-271-2008-00 047-C.

## Funding

This work was supported by an INBRE grant from the NIGMS (P20GM103446) to [C.J.C], a COBRE (5P20GM103653), an R21 from NINDS (R21NS109627) to [B.L.E.], and a James S. McDonnell Foundation grant to [B.L.E.]. Opinions are not necessarily those of the NIH.

## SI Appendix

We discuss how we reconstructed pathways for diffusion MR tractography, the tract-tracers we selected to study in mice, and how we varied imaging parameters to ensure our results were not driven by outliers.

### Diffusion MR Tractography

Diffusion MRI data were processed with Diffusion Toolkit (www.trackvis.org; threshold angle: 45 angles for 13/14 brains). In some panels, fibers were skipped for visualization but not for the analyses. With the exception of the human brain scans from the Allen Brain Atlas, we used high angular resolution diffusion MR imaging (HARDI) tractography to generate whole brain tractography. The orientation distribution functions (ODFs) were normalized according to the maximum ODF length within each voxel. Fractional anisotropy (FA) was calculated from orientation vectors by fitting the data to the tensor model (1). We used fiber assessment by continuous tracking (FACT) with HARDI. No fractional anisotropy threshold was applied in reconstructing tracts, which is consistent with previous work (1-4). We used TrackVis (http://trackvis.org) to visualize and quantify pathways. Table S9 provides details on the individuals scanned, including their age, and the spatial resolution used.

### Tract-Tracers

We compared diffusion MR tractography with tract-tracers in mice to assess which metrics should be extracted from tractography. We compared diffusion MR tractography in P60 mouse brains with viral tracer experiments, which involved EGFP injections into selected regions of the mouse brain. These data were made available by the Allen Mouse Brain Connectivity Atlas (https://connectivity.brain-map.org/; Fig. 4). We identified injection sites and set regions of interest (ROIs) to compare projections identified from the tract-tracers with those from tractography. We found strong concordance in the location and the orientation of pathways coursing through the white matter. We observed that tracts did not necessarily penetrate the grey matter in the same location as the tract-tracers. Axons make an abrupt turn at the junction of the grey and the white matter (Fig. 4). These sharp turns challenge the accuracy of tractography, as evident from the qualitative observations of tract-tracers and tractography (Fig. 4 and Fig. S11).

We compared tract-tracers with diffusion MR tractography to guide the development of quantitative approaches to study FC pathways. As there is no evidence of a relationship between fiber numbers and circuits (e.g., axons), we did not quantify fibers. Instead, we selected voxels through the FC white matter, classified pathway types, and quantified the proportion of pathway types across the FC and the PFC. In relatively rare cases, we refrained from classifying the pathway types if the tractography was not clearly classifiable.

### Varying Sampling and Imaging Procedures of Diffusion MR Scans

Variation in imaging resolution was inevitable considering the wide range of brain sizes used in the present study. We tested how sampling, resolution, and imaging protocols impact the percentage of pathway types, as it is unclear how these parameters impact results from tractography. We randomly sub-sampled the number of sites and extracted the average proportion of pathway types in the FC of each individual (Fig. S13). Variation in the percentage of pathway types was minor regardless of the number of randomly selected voxels, imaging procedures, and resolution (Fig. S12–S14). Our analyses suggest that the differences in the pathway types or a lack thereof are robust to sampling size, resolution, and imaging protocols.

### Transcriptional Variation Across Layers

We measured the intensity of gene expression across layers in a manner similar to that done previously (5-7). We downloaded ISH images of the FC from humans, macaques, and mice from the Allen Brain Atlas. We analyzed the expression of select SE genes within layers II–IV and V–VI. The boundary across layers IV and V was based on the cytoarchitecture from Nissl staining and *RORB* mRNA expression. Layer IV was defined as a cell-dense zone characterized by preferential expression of *RORB*. Layer VI was bound by the white matter. We measured the expression intensity from ISH images of *CRYM* in humans, macaques, and mice (Table S4). We used Image J software to randomly select areas within the FC, then we placed a rectangular grid to capture gene expression intensity across layers II–IV and V–VI. The grid was perpendicular to the cortical surface. The height varied with cortical thickness. Frame widths were 1,000 µm in mice, macaques, and humans. We binarized the images and measured the intensity of expression in layers II–IV and V–VI. We computed the ratio of these values to compare the relative expression of genes of interest in the upper and the lower layers. A value >1 indicates that the gene is preferentially expressed in layers II–IV.

**Fig. S1.**
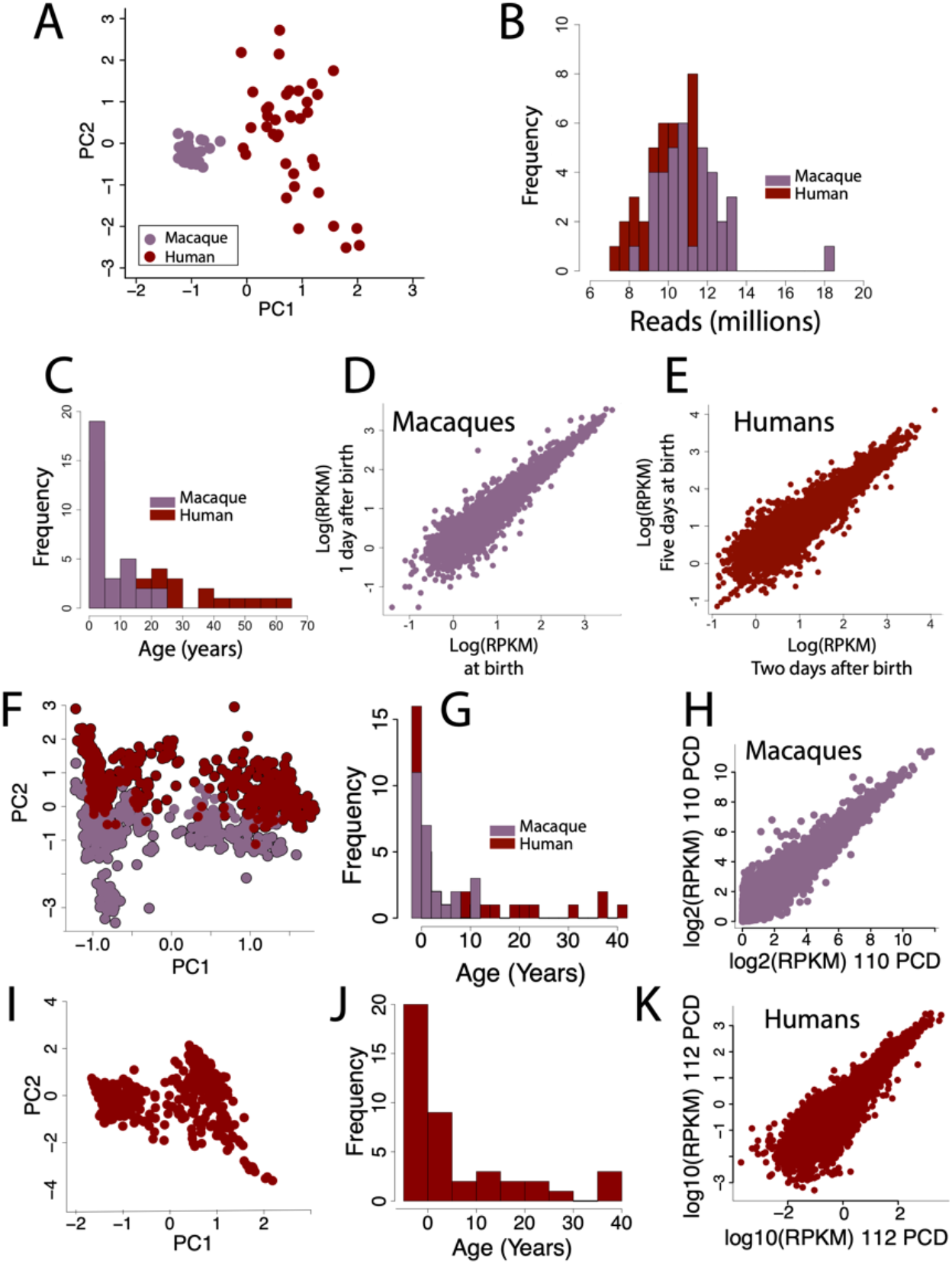
We used RNA sequencing datasets from Liu et al. (8) (A-E), Zhu et al. (9) (F-H), and the BrainSpan Atlas of the Developing Human Brain (10) (I-K). (C-E) We evaluated PCA on log-transformed data (A), reads (B), age ranges (C) and replicates (D-E) as exemplified for the Liu et al. (8), dataset to investigate potential outliers and quality of RNA sequencing across species and data-sets. (A, F, I) PCAs on log10-transformed reads per kilobase per million (RPKM) do not show any obvious outlier. Rather, samples cluster primarily by species. (C, G, J) Histograms show the distribution of ages of samples used in this study. The age of these samples, which are expressed in years after birth, overlap across the two species. (D, E, H, K) Scatter plots of log10-transformed RPKM values from biological replicates permit evaluating variance in gene expression from samples collected at very similar (D, E) or identical (H, K) ages. These plots show that variance decreases with increasing expression across replicates. Given these observations, we filtered genes based on expression levels to focus on genes that are relatively well and consistently expressed across replicates in our computational analyses.

**Fig. S2.**
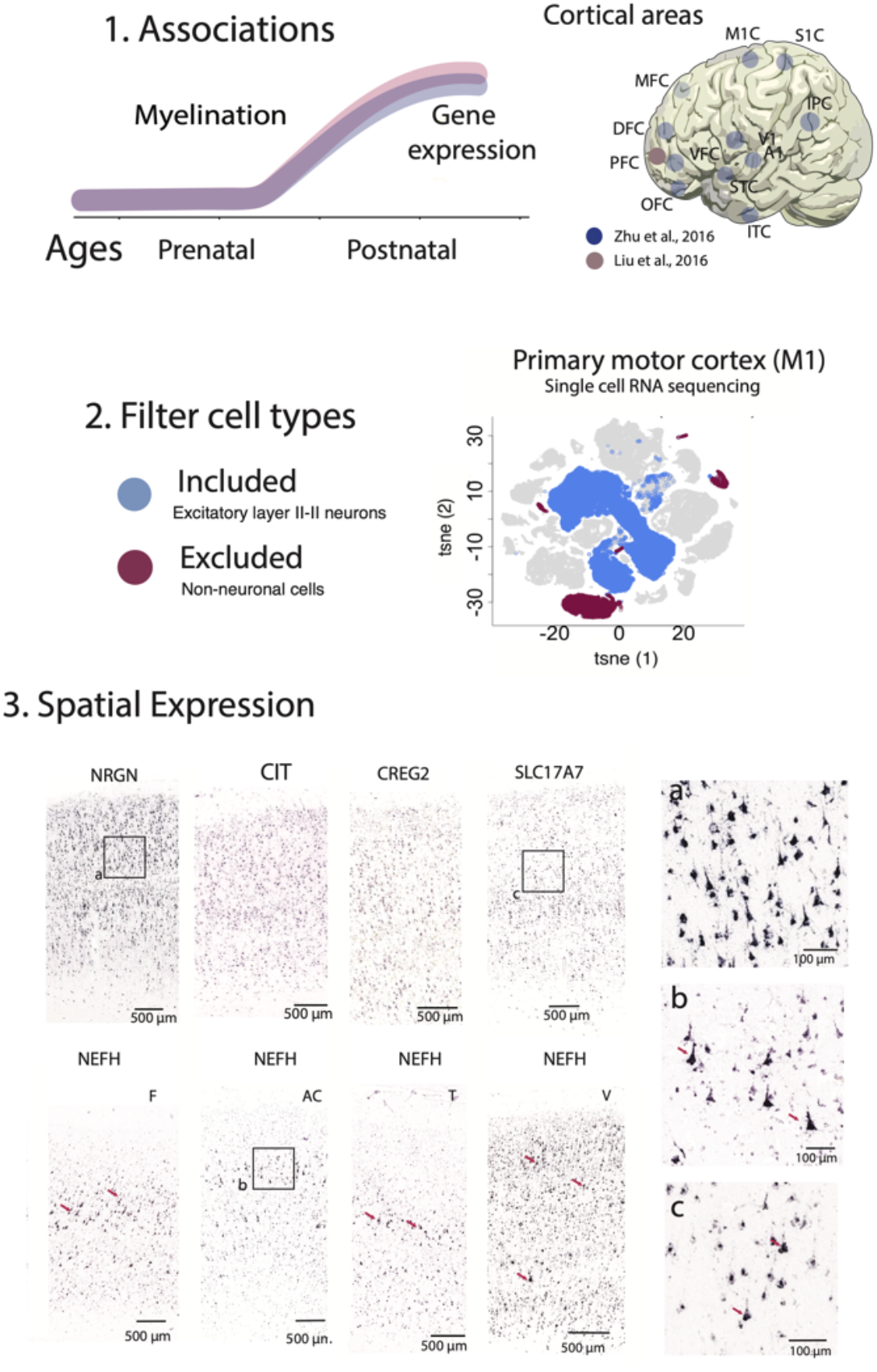
We summarize procedures to identify LRP markers and include in situ hybridization images to showcase the expression of select LRP across layers of the cortex. (1) We systematically tested for associations between myelin water fraction and gene expression across cortical areas. Only genes that systematically correlated with MWF and gene expression were considered candidate LRP markers. (2) We used single-cell RNA sequencing data from the primary motor cortex to consider only genes that are expressed by layer II-III neurons and exclude those expressed by non-neuronal cells (e.g., astrocytes, oligodendrocytes, microglia). We did not exclude neurons from other layers because cortico-cortical projections may emerge from neurons across layers (e.g., layers V-VI). (3) We used in situ hybridization made available from the Allen Brain Institute to evaluate spatial variation in transcriptional profiles of genes across the depth of the frontal cortex (top panel) and across cortical areas (lower panel) in humans at ages>20 years. We also evaluate the spatial expression of studied genes (e.g., *NRGN, CIT, CREG2*, and *SLC17A7*) to ensure that they are expressed by layer II-III neurons. Close-up views from these panels (*NRGN*: a *NEFH*: b; *SC17A7*: c) show that these genes are expressed in large pyramidal neurons of layers II-III and that their expression spans the depth of the cortex. These genes are also systematically observed to be expressed by pyramidal neurons across the cortex. Shown here is *NEFH* expression by large pyramidal neurons across the frontal (F), anterior cingulate (AC), temporal (T), and visual (V) cortex. The observation that these select LRP markers are expressed in large layer III pyramidal neurons, and that LRP markers include *NEFH*, which is a well-known marker of LRP neurons, supports the validity of our analyses in transcriptionally defining LRP neurons. These observations support the observation that our computational approaches identify LRP markers.

**Fig. S3.**
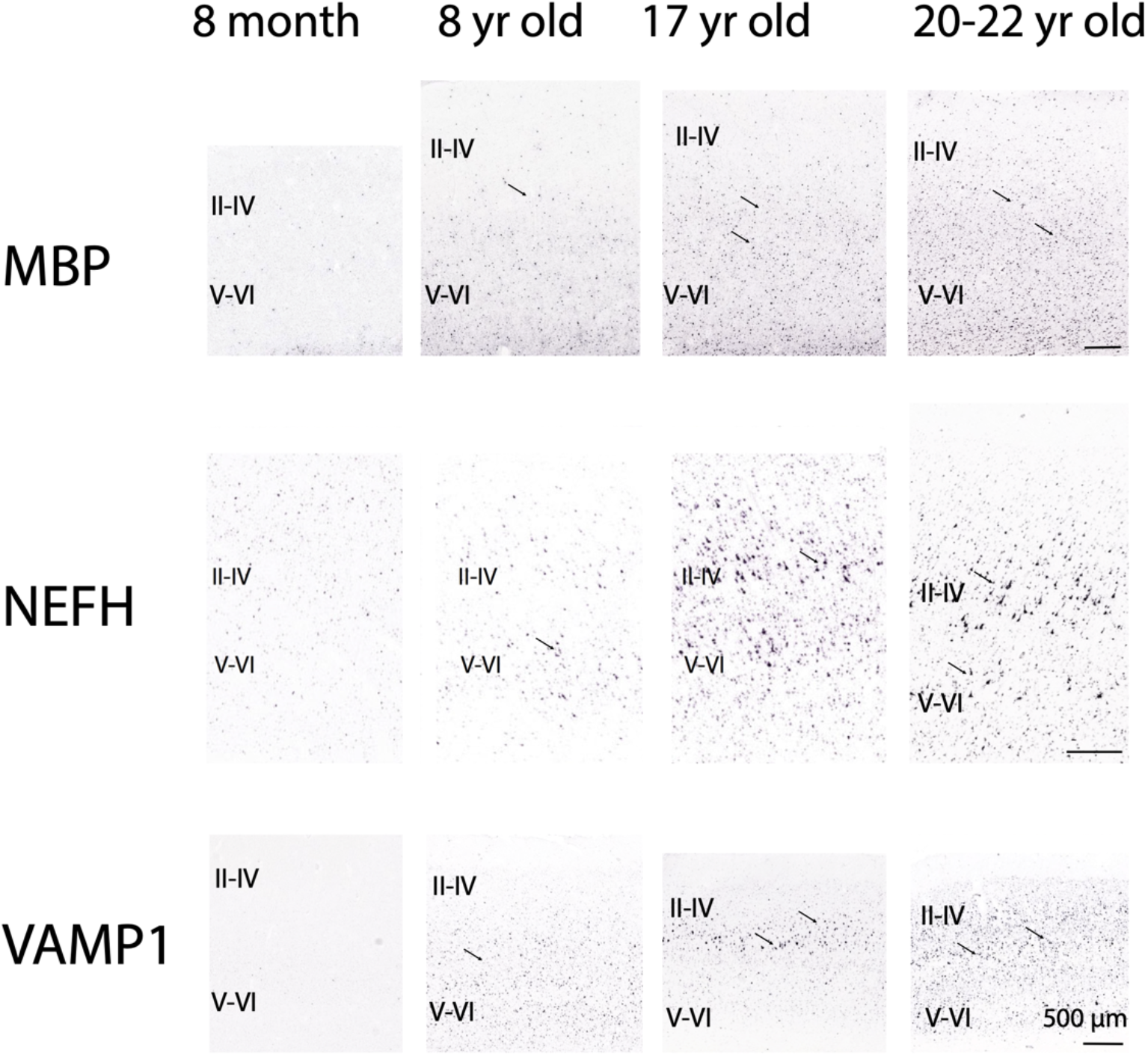
Spatiotemporal transcriptional profiles of myelin basic protein (*MBP*) and select LRP markers (*NEFH* and *VAMP1*) in the human cortex show that the expression of these genes increases postnatally. Myelination according to *MBP* expression is weak shortly after birth but systematically increases during postnatal development. A similar situation is observed for *NEFH* and *VAMP1*, which are weakly expressed in the cortex shortly after birth but systematically increase towards adulthood. Arrows point to large pyramidal neurons. These qualitative observations align with our computational analyses. That is, expression of identified LRP markers correlates with myelination and systematically increases during postnatal development. In situ hybridization images are from the Allen Brain Institute. Expression of *NEFH* and *MBP* in the frontal cortex and *VAMP1* in the visual cortex is shown.

**Fig. S4.**
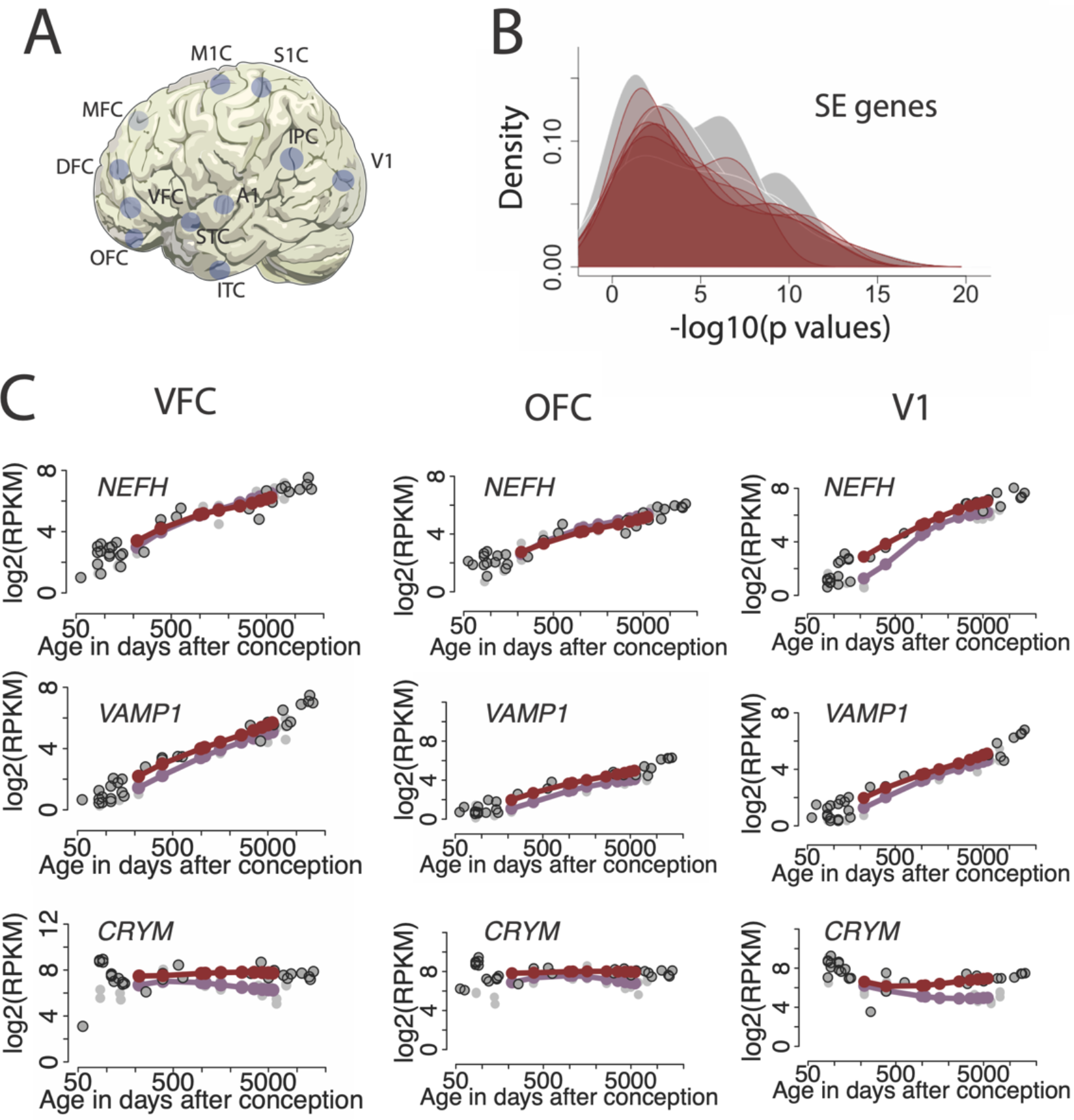
Temporal variation in supragranular-enriched (SE) genes in humans and macaques are very similar in the two species once cross-species variation in developmental schedules are aligned. Data are from Zhu et al. (9), We extracted normalized gene expression at corresponding time points across cortical areas (*A*) and correlated normalized gene expression across species. Here, age in macaque is translated to humans according to Fig. 1. (*B*) Significance tests overlap extensively across areas, which indicate temporal profiles in SE genes are very similar in humans and macaques. (*C*) Indeed, temporal profiles in SE gene expression (e.g., *NEFH, VAMP1, CRYM*), from the orbitofrontal cortex (OFC), ventral frontal cortex (VFC), and primary visual cortex (V1) appear highly similar in humans and macaques. Together, these data showcase a lack of evidence for protracted development in FC circuits in humans relative to macaques. Additional abbreviations: A1: primary auditory cortex; DFC: dorsolateral frontal cortex; IPC: inferior parietal cortex; ITC: inferior temporal cortex; MFC: medial frontal cortex; M1C: primary motor cortex; S1C: primary somatosensory cortex; VFC: ventral frontal cortex.

**Fig. S5.**
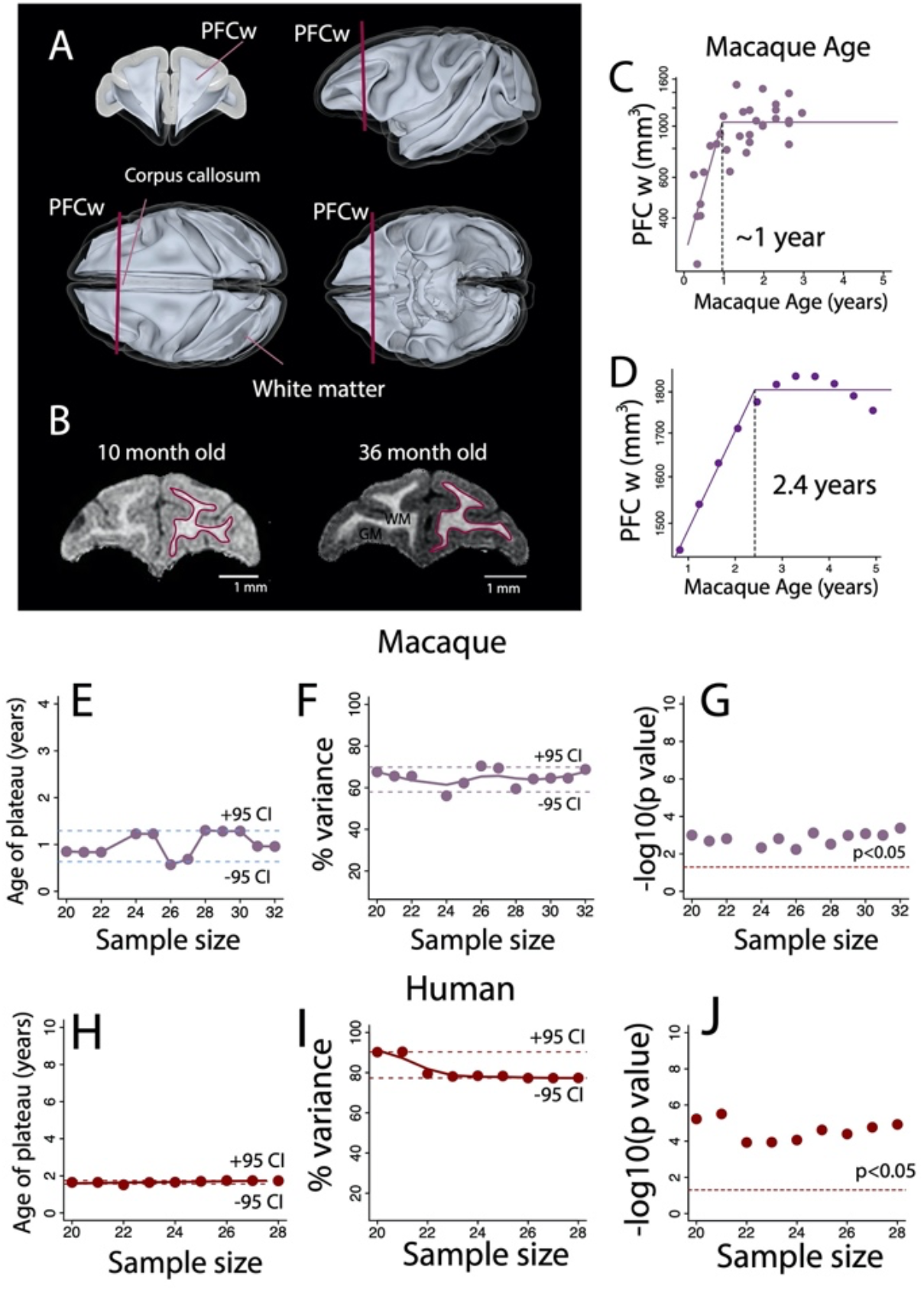
We tested whether age of growth cessation is driven by outliers and is consistent across studies. To do this, we evaluated the impact of subsampling on the age of growth cessation. We also compared our determination of PFC white matter growth with those collected from prior studies to ensure the reproducibility of our results. (*A*) The PFC white matter (PFCw) was defined as the white matter anterior to the corpus callosum as exemplified on a macaque white matter brain template (11, 12). Red vertical bars illustrate the posterior boundary of the PFCw. (*B*) Red contours label the boundary between the grey (GM) and white matter (WM) in the coronal plane of a 10-month and 36-month-old macaque brain to show PFCw delineations. (*C* and *D*) The growth trajectories of the PFCw are similar across studies. We fit a nonlinear regression (easy model 3) with the log-transformed PFCw volume regressed against age expressed in days after conception. The PFCw ceases to grow approximately 1 year after birth in macaques. We applied the same nonlinear regression on other data (13) to assess the range of variation in the timetable of PFCw growth (Fig. 1). We first fit a smooth spline with the PFCw volume versus age for males and another one for females. We subsequently fit another smooth spline through these data to average values across males and females. To test how sampling impacts the age of growth cessation, we subsampled the number of individuals and applied a non-linear regression on the log-transformed PFCw volumes versus age expressed in days after conception in macaques and humans (*E*–*J*). These nonlinear regressions capture when the growth of the PFCw ceases (*E*–*G*). We extracted the age in which PFCw growth ceases according to these nonlinear regressions (*E* and *H*), the percentage of variance accounted for by the model (*F* and *I*), and significance tests (i.e., p values)(*G* and *J*). (*E*) The 95% confidence intervals show that the macaque PFCw ceases to grow between ∼0.6 and ∼1.3 years of age. (*F* and *I*) These nonlinear regressions systematically account for a high percentage of the variance (∼>70%) and (*G* and *J*) are statistically significant (p<0.05). These analyses, along with those in Fig. 1, showcase a lack of evidence for protracted PFC development in humans.

**Fig. S6.**
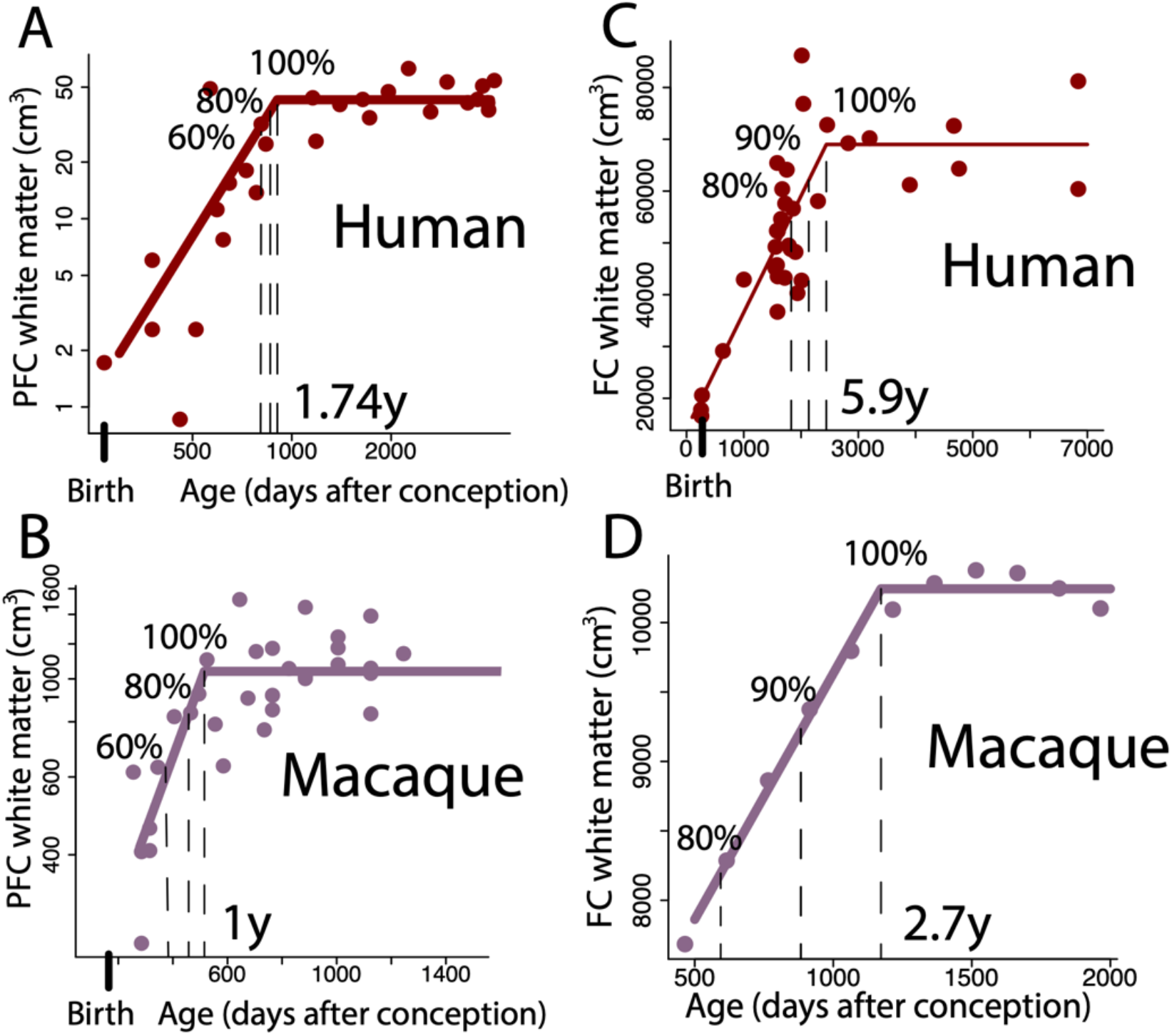
We extracted time points from the FC and PFC growth trajectories in humans and macaques and tested whether subsampling impacts the age of growth cessation. We fit a non-linear regression (easynls, model=3) through the FC white matter volume and the log10 PFC white matter regressed against age in humans and macaques. We extracted the age at which the PFC and FC white matter reach a particular percentage of adult volumes. The choice of epochs were based on the age ranges from which volumetric data were available. These non-linear regressions show that the PFC white matter ceases to grow at 1.74 years of age in humans (*A*) and 1 year of age in macaques (*B*). The FC ceases to grow at 5.9 years of age in humans (*C*) and 2.7 years of age in macaques (*D*). The growth of the white matter may certainly extend beyond these ages but at a slower rate beyond the identified ages.

**Fig. S7.**
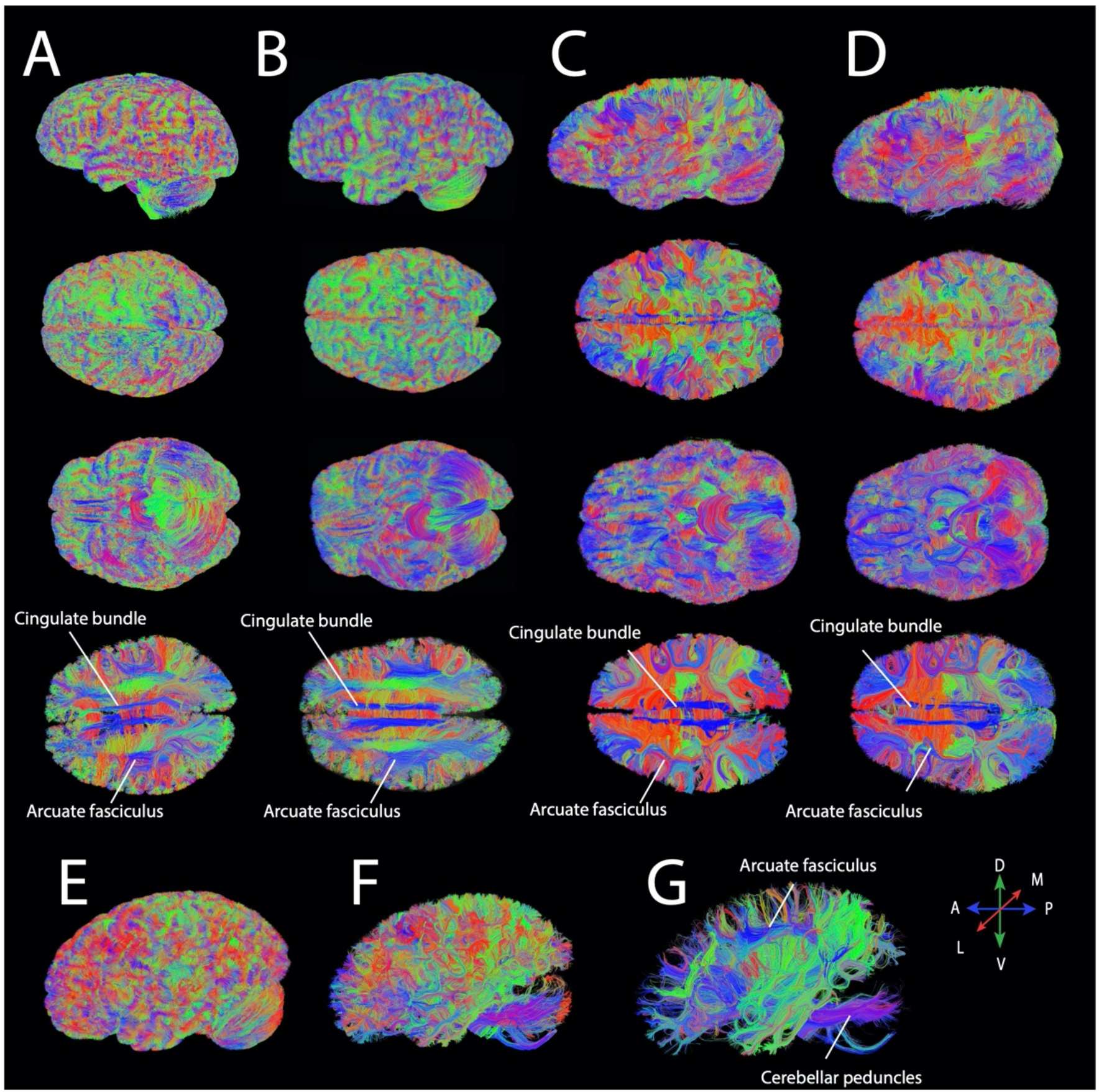
Some examples of the diffusion MR tractography of human brains in lateral, dorsal, and ventral views. The scans used in the study show consistency across individuals and imaging protocols. Whole brains (*A*–*D*) and a left hemisphere (*E*–*G*) were scanned on a 3T Siemens Tim Trio scanner at the Massachusetts General Hospital Athinoula A. Martinos Center for Biomedical Imaging, some of which are also housed by the Allen Brain Institute (*C* and *D*). A 1-mm thick horizontal slice filter set through the human brains show fibers coursing across the brain such as the cingulate bundle and the arcuate fasciculus. These data show consistent high resolution in fiber tracking, and consistency across individuals and scanning protocols. (*C* and *D*) Two of the brains used from the Allen Brain Atlas. These scans vary in their resolution (isotropic: 1.2 mm and 0.9 mm, respectively). (*E*–*G*) A left hemisphere shown at different minimum length thresholds (E: 0 mm, F: 20 mm, and G: 50 mm). We visualized pathways at different thresholds to reveal fibers of various lengths coursing through the white matter such as the (G) arcuate fasciculus. The color-coding of tractography was based on a standard red–green–blue (RGB) code based on average fiber direction. Abbreviations include: A: anterior; P: posterior; R: rostral; C: caudal; M: medial; L: lateral; D: dorsal; and V: ventral.

**Fig. S8.**
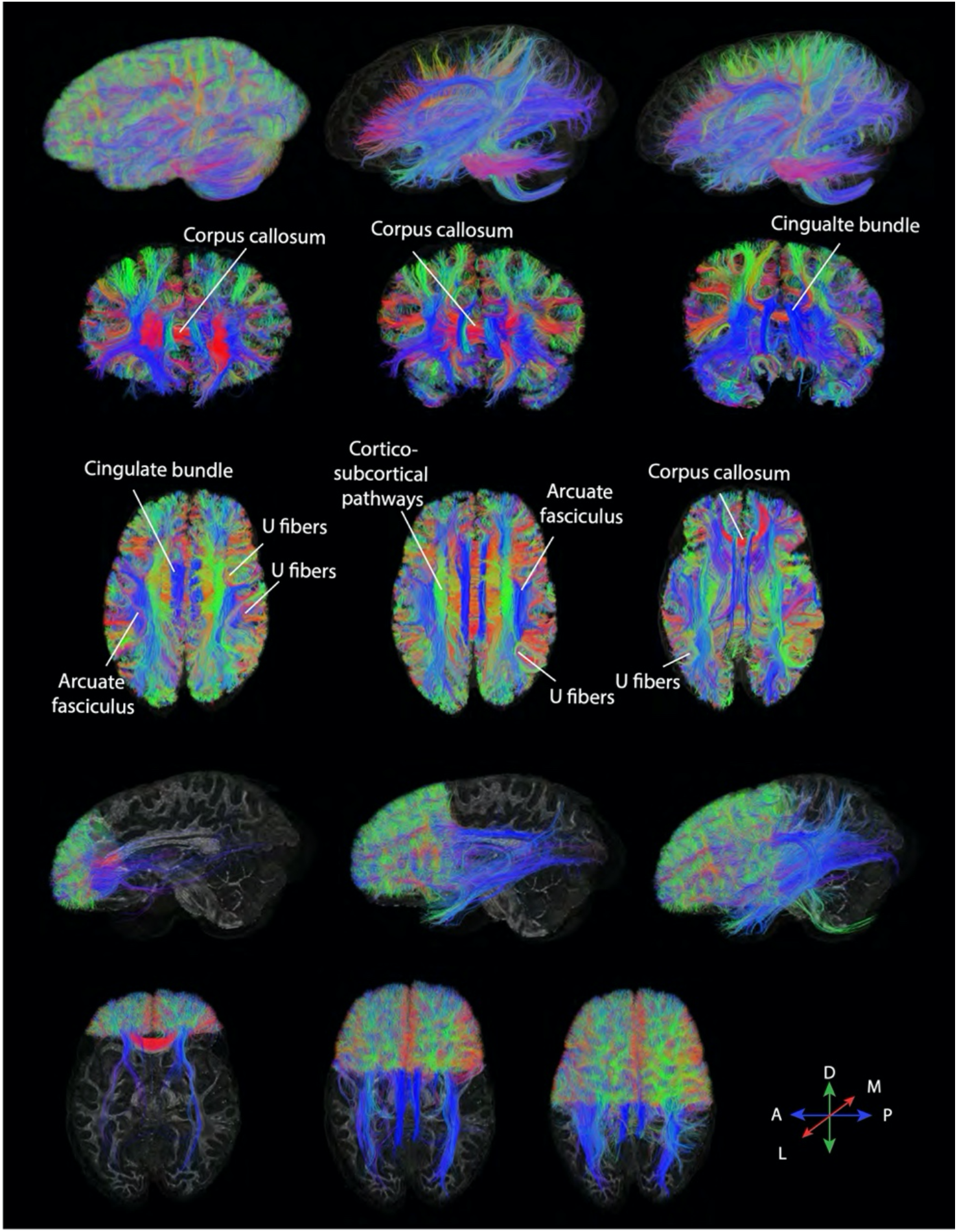
Whole brain tractography through the human brain with variable minimum length thresholds highlight various pathways (0 mm, 40 mm, 20 mm). These thresholding parameters reveal pathways coursing across the anterior to posterior axis, and also the presence of relatively few long pathways (greater than 40 mm) coursing through the FC. In other words, there appears to be preponderance of short fibers within the FC. Coronal slices (width: 1 mm) show fibers coursing in various directions through the FC. The human FC is composed of a number of cross-cortically projecting fibers. Horizontal slices (width: 1 mm) show well-known pathways coursing across cortical white matter, some of which span the FC (e.g., cingulate cortex and corpus callosum). Coronal and horizontal slices (second and third from the top) of various sizes show FC fibers. In these panels, 90% of tracked fibers are skipped to more easily visualize fibers coursing through the human FC. A number of relatively short and long-range fibers are observed coursing across the FC white matter. Taken together, these observations suggest that the FC is largely composed of pathways projecting within the FC. For visualization purposes, the color-coding of tractography was based on a standard red–green–blue (RGB) code based on fiber direction. Abbreviations include: A: anterior; P: posterior; R: rostral; C: caudal; M: medial; L: lateral; D: dorsal; and V: ventral.

**Fig. S9.**
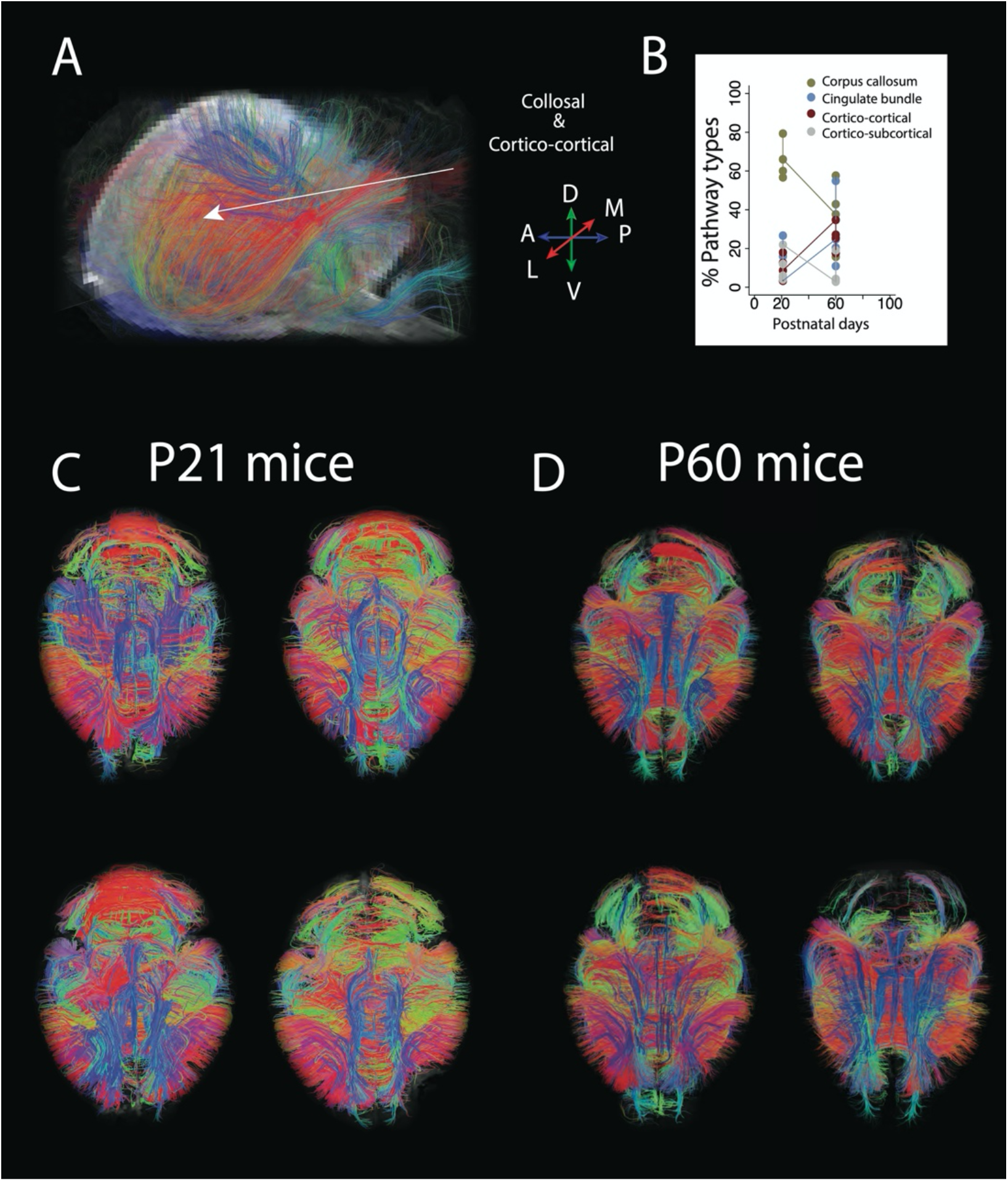
Diffusion MR tractography of mouse brains at P21 and P60 reconstructed with HARDI. These post-mortem mouse brains were scanned with the 9.4T scanner at the University of Delaware. (*A*) Close up views through anterior regions of the mouse cortex show fibers coursing across the medial to lateral and anterior to posterior axes of the cortex. The fibers coursing across the medial to lateral axes should represent callosal fibers and pathways connecting cortical and subcortical structures. Fibers coursing across the anterior to posterior axis include the cingulate bundle as well as other anterior to posterior cross-cortical pathways. The tractography is very similar across individuals of the same and different ages. (*B)* We quantified pathway types in the brains of P21 and P60 mice. Although the corpus callosum is relatively expanded in P60 versus P21 mice, a factor for age is not significant when an ANOVA is performed on these data. Dorsal views of whole brain tractography of mouse brains at (*C*) P21 and (*D*) P60 show strong concordance across individuals and ages. The color-coding of tractography is based on a standard red–green–blue (RGB) code based on fiber direction. Abbreviations include: A: anterior; P: posterior; R: rostral; C: caudal; M: medial; L: lateral; D: dorsal; and V: ventral.

**Fig. S10.**
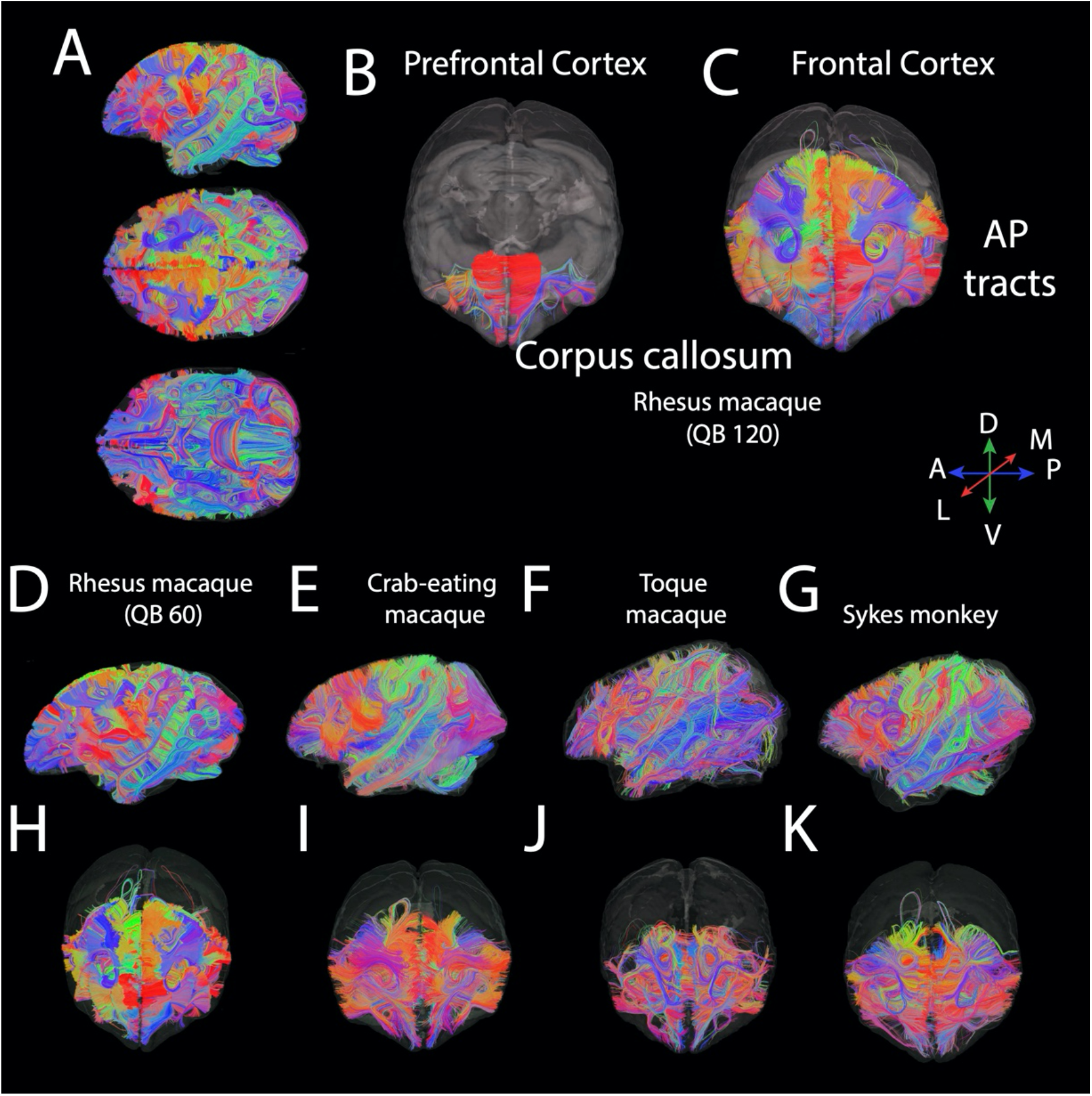
Brain tractography shows pathways coursing through the brain of Old World monkeys used in the present study. Those include several macaques such as a rhesus macaque (*A*–*C*; scanned with 120 directions, *D*: scanned with 60 directions), a crab-eating macaque (*E*), a toque macaque (*F*), and a Sykes monkey (*G*). The tractography is consistent across individuals. Slices set through the prefrontal cortex (*B*) and larger areas through the FC (*C* and *H–K*) capture fibers emerging from and terminating within the FC. These settings show that the FC is composed by a preponderance of pathways emerging and terminating within the FC. A minimum length threshold (15 mm) was set for visualizations purposes only. Brain masks are overlaid on these pathways. The color-coding of tractography is based on a standard red–green–blue (RGB) code based on fiber direction. Abbreviations include: A: anterior; P: posterior; R: rostral; C: caudal; M: medial; L: lateral; D: dorsal; and V: ventral.

**Fig. S11.**
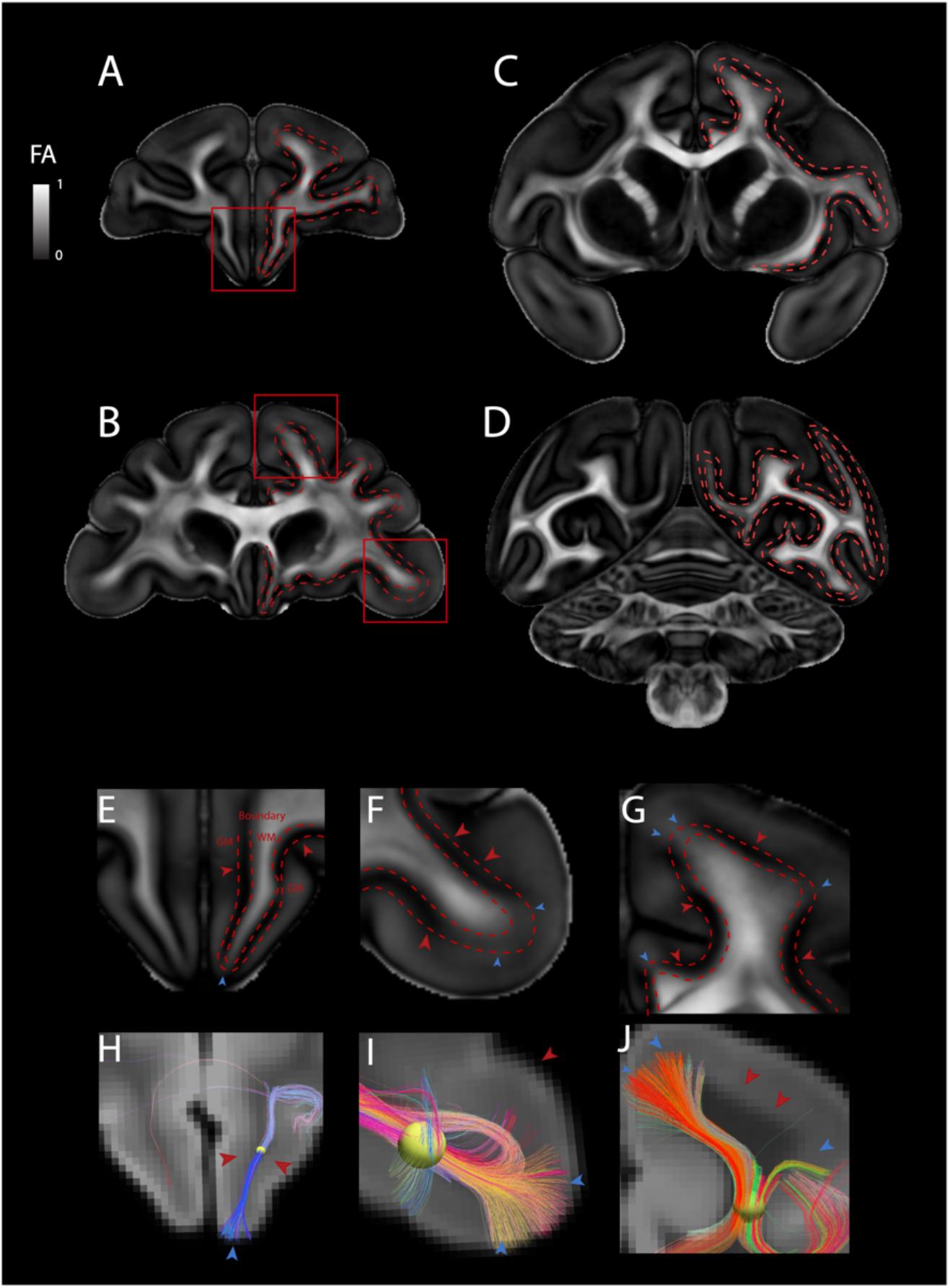
We refrain from considering fiber endpoints within the grey matter as a basis with which to classify pathway types. We showcase one example of biases in tractography across the macaque cortex, which could impact pathway classification. We considered fractional anisotropy (FA) images and tractography to identify possible sources of errors in tracking fibers at the grey to white matter boundary. For the most part, there is very low FA at the grey to white matter boundary, which means uncertainty in tracking fibers penetrating the grey matter. (*A*–*D*) Dashed lines at the intersection between the grey (GM) and white matter (WM) show FA is particularly low (dark) at this boundary. FA varies across the cortex, with high FA within gyri (that is, high likelihood of fiber crossing) and low FA towards sulci (low likelihood of fiber crossing). Spheres capturing fibers through the white matter show that fibers preferably penetrate the cortex within gyri. (*H–J*) Close up views through select cortical areas show that fibers preferentially penetrate the grey matter at areas of high FA (*E*–*J*, blue arrowheads) rather than low FA (red arrowheads). These observations suggest that fibers penetrating gyri may be over-represented at the expense of those closer to sulci because of increased uncertainty in tracking fibers penetrating the grey matter in zones within sulci. It is because of uncertainties in the tractography such as these that the classification of pathways are not constrained by their precise termination of fibers within the grey matter but focus instead on their orientation within the white matter. FA images and tractography are from Calabrese et al. (12).

**Fig. S12.**
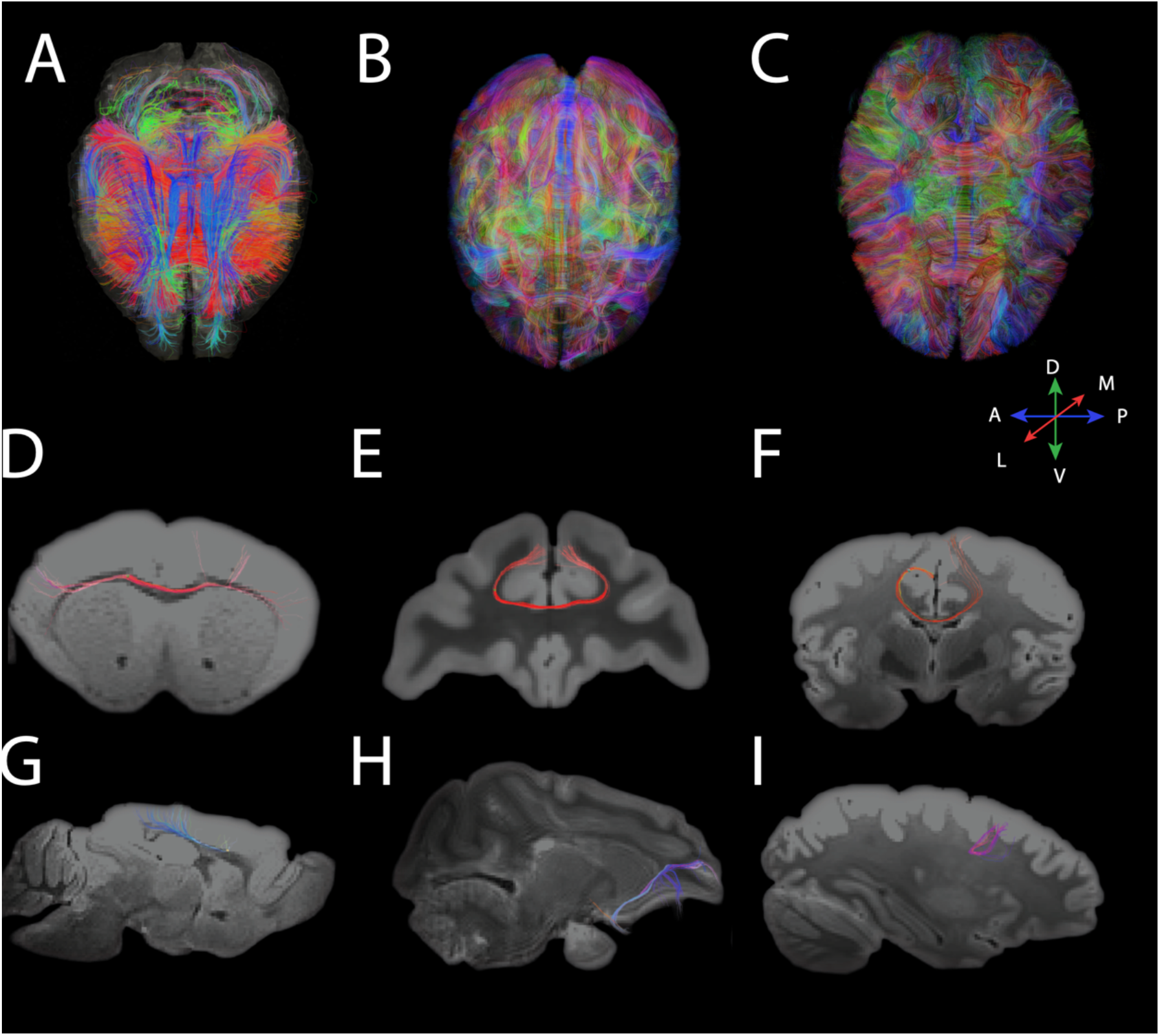
Tractography of fibers highlighting pathways coursing through the FC in mice (*A*), macaques (*B*), and humans (*C*). Brain pathways for macaques (*B*) and humans (*C*) are partially transparent. We randomly selected voxels to capture the relative percentage of pathways coursing through the FC of each to quantify the composition of the FC white matter across the three species. (*D*–*I*) Examples of pathways identified from randomly selected voxels include collosal fibers, U fibers, and long-range cortically-projecting pathways in mice (*D* and *G*), macaques (*E* and *H*), and humans (*F* and *I*).

**Fig. S13.**
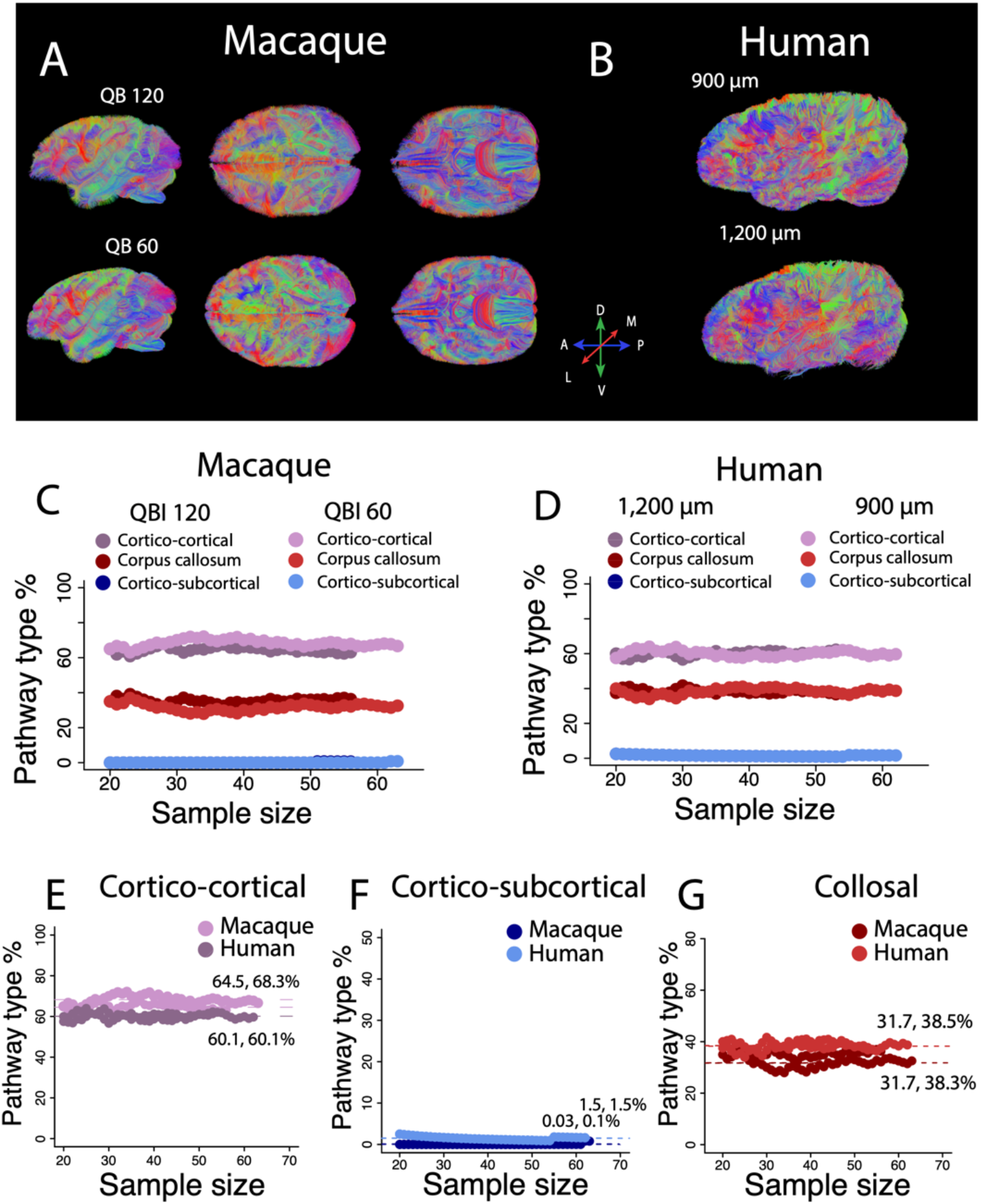
We considered how sampling and imaging parameters impact the proportion of pathway types in macaques and humans. Some brains scanned with different parameters are shown in *A and B*. We assessed the percentage of pathways through the FC white matter with varying (*A*) direction number (60 versus 120) and (*B*) resolution (900 µm versus 1200 µm). (*C*–*G*) The percentage of pathway types varies little with sample size in macaques (*C*) and in humans (*D*). Horizontal dashed lines and associated values show the relative mean pathway types per scanning parameters with varying sample size in humans and macaques. The percentage of cortico-cortical (*E*), cortico-subcortical (*F*), and collosal pathways (*G*) are similar in humans and macaques regardless of sampled voxels used to classify pathways. The percentage of pathway types are robust to variation in resolution and sampling. A minimum length threshold was set to 7.2 mm for humans and 5.8 mm for macaques for visualization purposes. This human brain is made available by the Allen Brain Institute (14). A: Anterior; P: posterior; M: medial; L: lateral; D: dorsal: V: ventral. In *A*, 90% of the fibers are skipped to better visualize pathways coursing through the macaque brain.

**Fig. S14.**
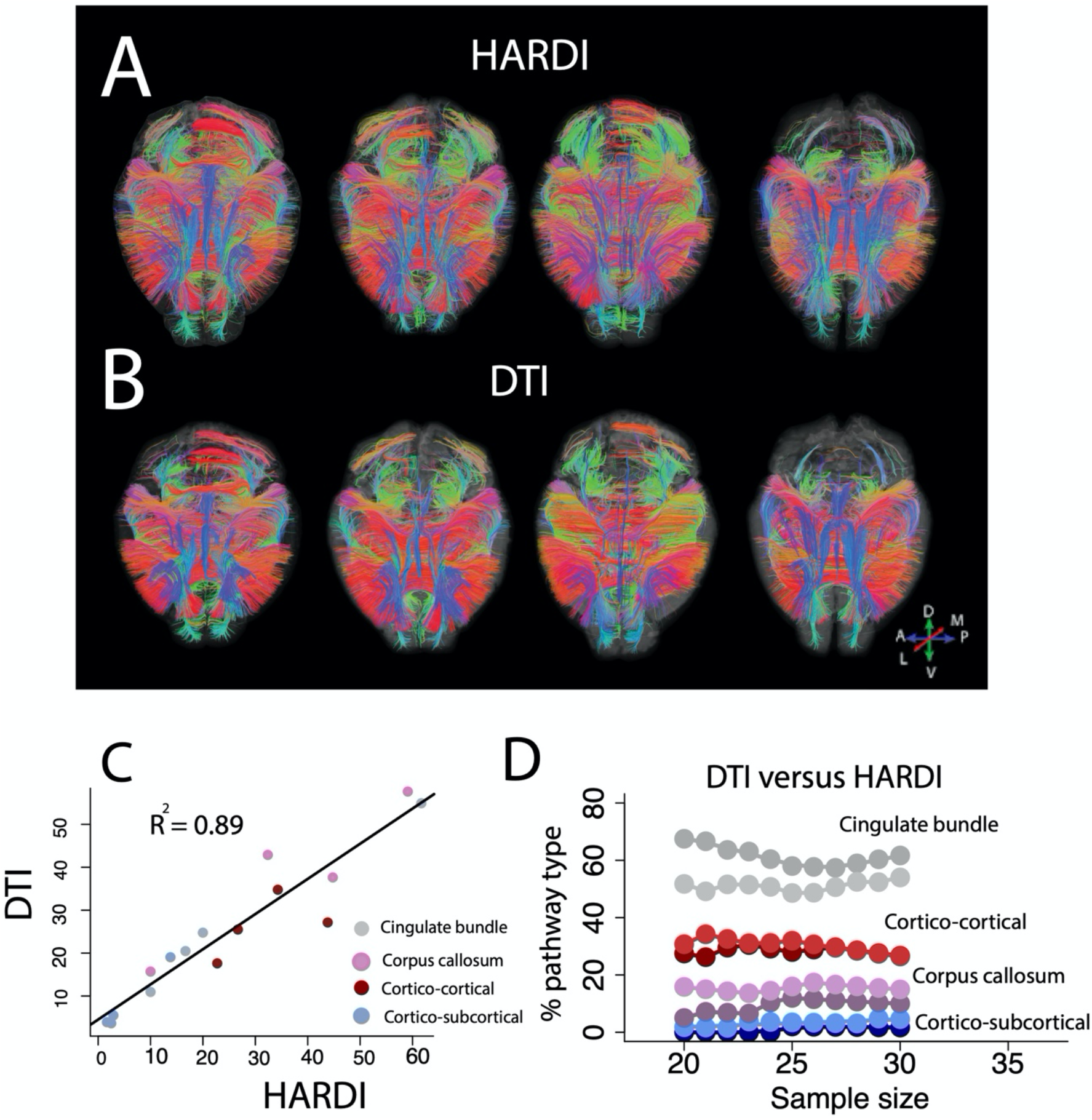
We tested how diffusion tensor imaging (DTI) and high angular resolution diffusion imaging (HARDI) impact pathway types coursing through the mouse FC white matter. We randomly selected voxels through the white matter and we quantified the percentage of pathway types reconstructed with DTI and HARDI. (*A* and *B*) Dorsal views of tracts reconstructed with HARDI. (*A*) appear similar to those reconstructed with DTI (*B*). (*C*) The relative percentages of pathway types reconstructed at P60 with DTI strongly correlate with those with HARDI (R2=0.89; p<0.01, n=16). (*D*) We randomly subsampled voxels and computed the relative percentages of pathways coursing through the mouse FC cortex to determine whether these percentages are relatively invariant with respect to variation in tractography and sample size. These analyses suggest that the relative percentages of pathways withstand variation in parameters and sampling strategy. In *A* and *B*, we selected fibers with a 2 mm minimum length threshold for visualization purposes only. Abbreviations: A: anterior; P: posterior; R: rostral; C: caudal; M: medial; L: lateral; D: dorsal; and V: ventral.

**Fig. S15.**
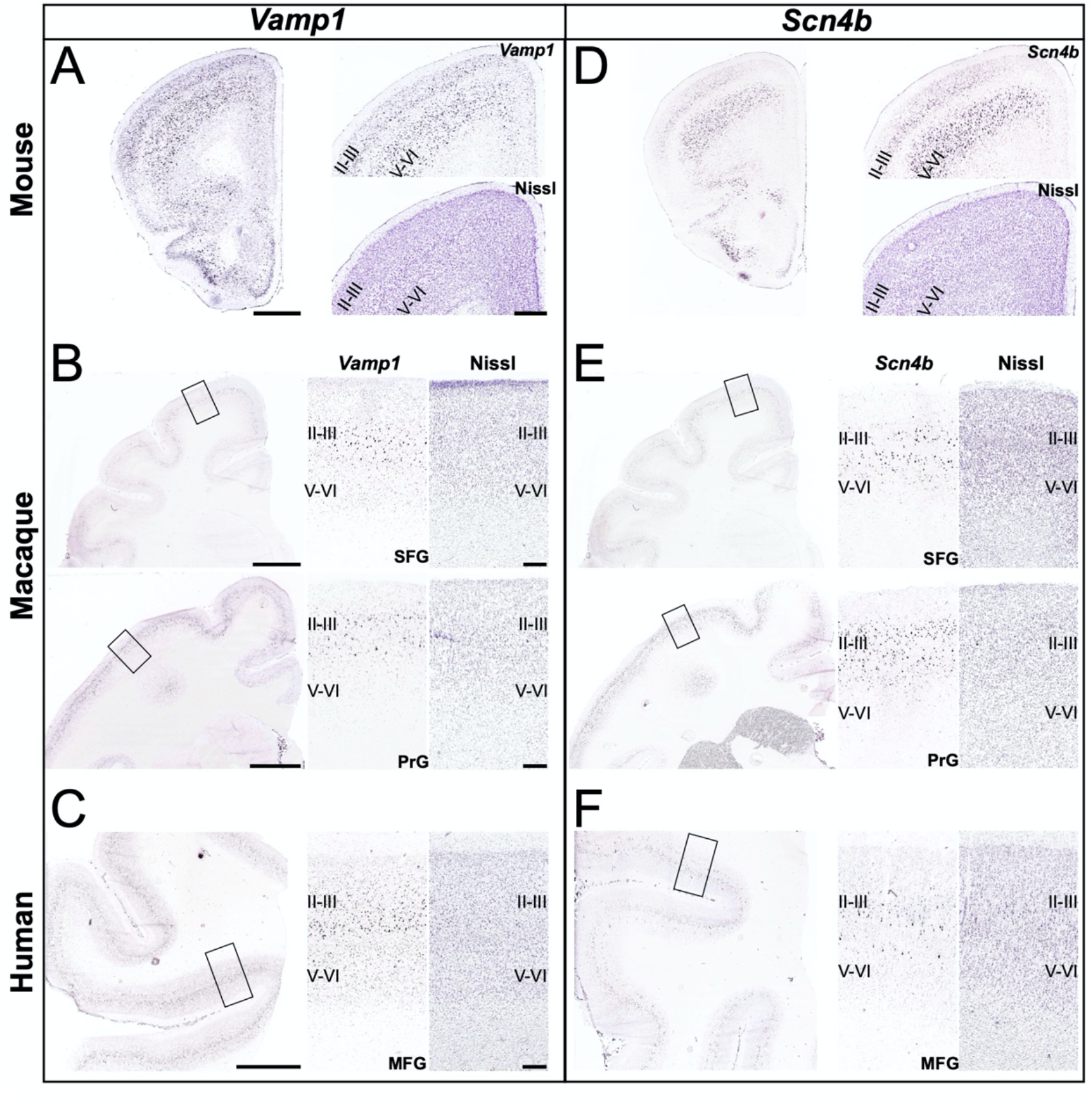
Expression of the SE genes *Vamp1* and *Scn4b* in the FC of mice, macaques, and humans. *In situ* hybridization and corresponding Nissl stains are from the Allen Institute for Brain Science. *(A–C) Vamp1* and (*D–E*) *Scn4b* expressions in coronal sections through the FC of (*A* and *D*) a mouse at P56 (male), (*B* and *E*), a macaque at 48 m (male), and (*C* and *F*) an adult human (46-year-old male control). (*A*) *Vamp1* and (*D*) *Scn4b* expressions in the mouse FC cortex region are shown, along with enlarged views and corresponding Nissl images. Both genes appear to be widely expressed throughout layers, including layers II-III. In macaques, the ISH signal was variable (n=3). However, staining was observed in a region anterior to and within the motor cortex. Sections through these regions in the macaque, along with enlarged detail views, show (*B*) *VAMP1* and (*E*) *SCN4B* expression in layers II-IV of the superior frontal gyrus (SFG) and precentral gyrus (PrG). The ISH signal was also variable in humans in the dorsolateral PFC (DL-PFC). A section through the middle frontal gyrus (MFG) in the DL-PFC shows that (*C*) *VAMP1* and (*F*) *SCN4B* is expressed in layers II-III, similar to what is observed in macaques. In macaques and humans, *VAMP1* and *SCN4B* appear preferentially expressed in layers II-III. This is in contrast to mice, in which they are more widely expressed across layers. All *in situ* hybridization and Nissl images were obtained from the Allen Institute for Brain Science. Scale bars: A, 1 mm and 500 µm; B, 5 mm and 500 µm; C, 5 mm and 500 µm.

**Fig. S16.**
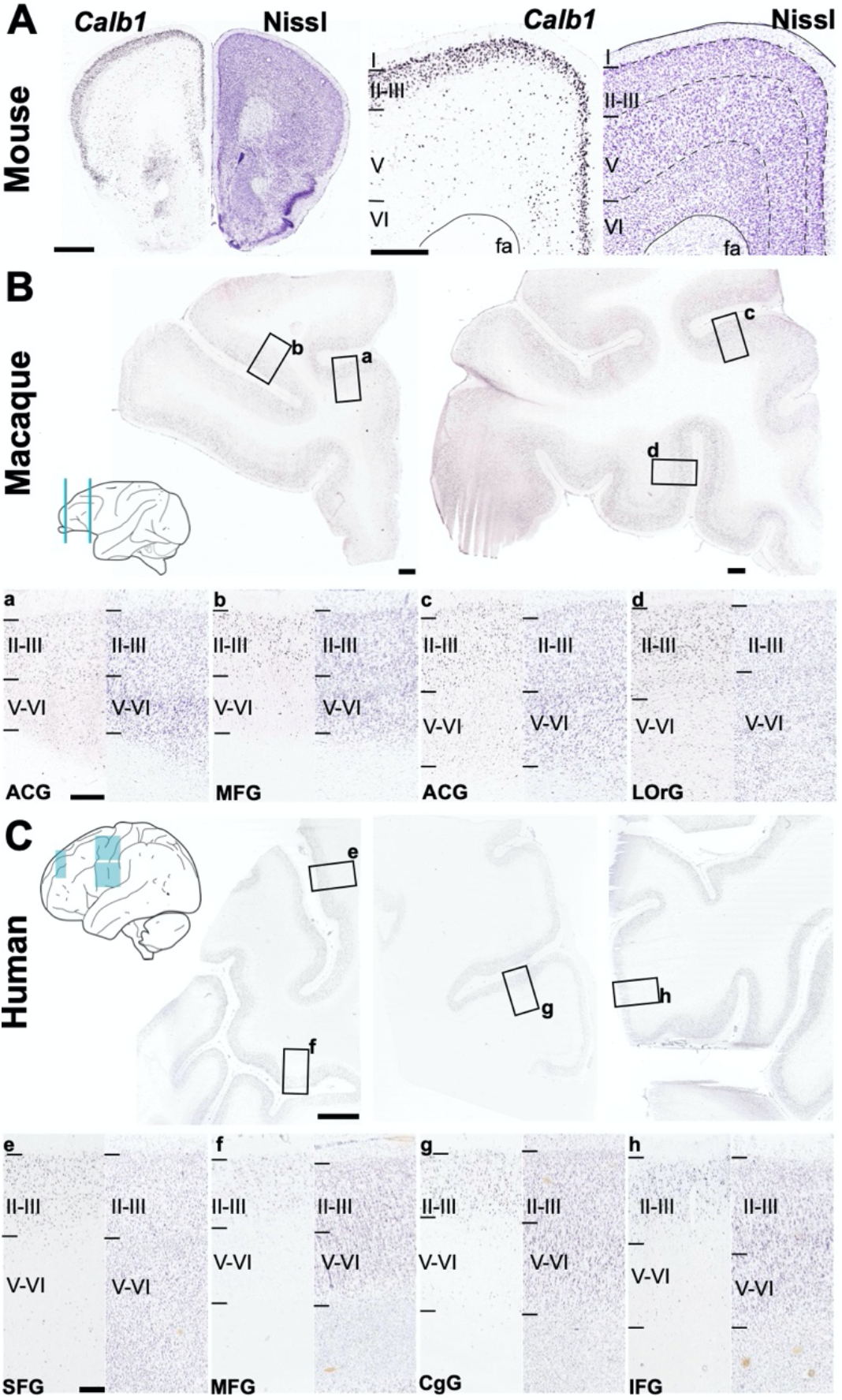
Expression of *CALB1* in cortical layers II-III of the FC across mice, macaques, and humans. (A) Coronal sections through a P56 mouse cortex show that *Calb1* is restricted to layers II-III. The enlarged detailed views show a region of the primary and secondary motor cortex. The anterior forceps of the corpus callosum (fa) is also shown. *Calb1* expression in the FC of (*B*) macaques (n=3) and (*C*) humans (n=15) is similar and shows the expansion of cortical layers II-III in the two species. (*B*) Sections through two regions of the FC (blue lines, schematic) of a macaque (48-m, male) show *Calb1* expression and enlarged detailed views, with corresponding Nissl images, of several gyri: *a* and *c*, anterior cingulate gyrus (ACG); *b*, middle frontal gyrus (MFG); *d*, lateral orbital gyrus (LOrG). (*C*) Sections and enlarged detailed views (with corresponding Nissl images) of three regions of the FC in a human (28-year-old male control) (blue boxes, schematic) showing *Calb1* expression in the following gyri: *e*, superior frontal gyrus (SFG); *f*, middle frontal gyrus (MFG); *g*, cingulate gyrus (CgG); and *h*, inferior frontal gyrus (IFG). All *in situ* hybridization and Nissl images were obtained from the Allen Institute for Brain Science. Schematics modified from the NIH Blueprint Non-Human Primate (NHP) Atlas and the Allen Human Brain Atlas. Scale bars: A, 1 mm and 500 µm; B, 1 mm and 500 µm; C, 5 mm and 500 µm.

**Fig. S17.**
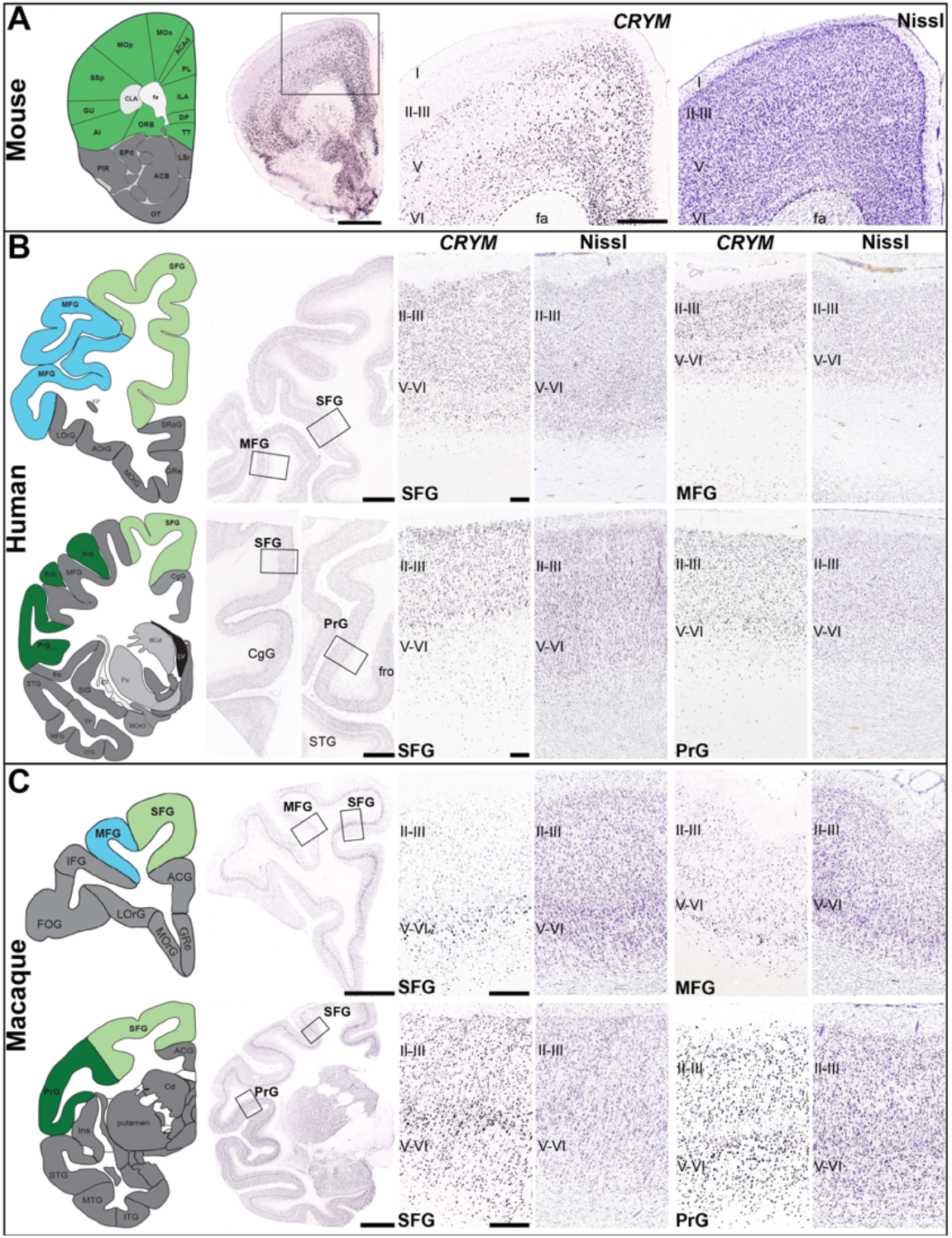
Variation in *CRYM* expression in the supragranular layers across the mouse, human, and macaque FC. *CRYM* in situ hybridization and Nissl data and modified atlas schematics are from the Allen Institute for Brain Science. (*A*) In the mouse (P56, male), *Crym* is not expressed in the supragranular layers (II-III) of the cortex. Enlarged detailed views show the primary and secondary motor areas of the cortex. For these views, a corresponding Nissl image is also shown. (*B*) By contrast, strong *CRYM* expression is observed in these cortical layers in humans (n=4), as observed in sections of several gyri at different rostral–caudal levels of the FC (31-year-old male control). Enlarged detailed views show *CRYM* expression and corresponding Nissl staining in the superior frontal gyrus (SFG) and middle frontal gyrus (MFG), and the SFG and PrG (precentral gyrus) in a rostral and caudal region of the FC, respectively. Overall, *CRY*M expression is similar across the four human specimens analyzed. (*C*) *CRYM* is also expressed in the supragranular layers in macaques (n=3), similar to humans, although its expression is variable across cortical regions. In the specimen shown (48 m, male), for example, *CRYM* expression in the SFG and MFG in a rostral region of the FC appears to be weaker than in similar gyri in more caudal regions. This variation is consistent across the three specimens analyzed. Abbreviations: CgG, cingulate gyrus; fro, frontal operculum; fa, anterior forceps of the corpus callosum; MFG, middle frontal gyrus (blue in schematic); PrG, precentral gyrus (dark green); SFG, superior frontal gyrus (light green); STG, superior temporal gyrus. For others, we used legends available at the Allen Brain Map (https://portal.brain-map.org/). Anatomic reference atlases used: the Allen Mouse Brain Atlas, the NIH Blueprint Non-Human Primate (NHP) Atlas, and the Allen Human Brain Atlas. Scale bars: A, 1 mm and 500 µm; B, 5 mm and 500 µm; C, 5 mm and 500 µm.

**Fig. S18.**
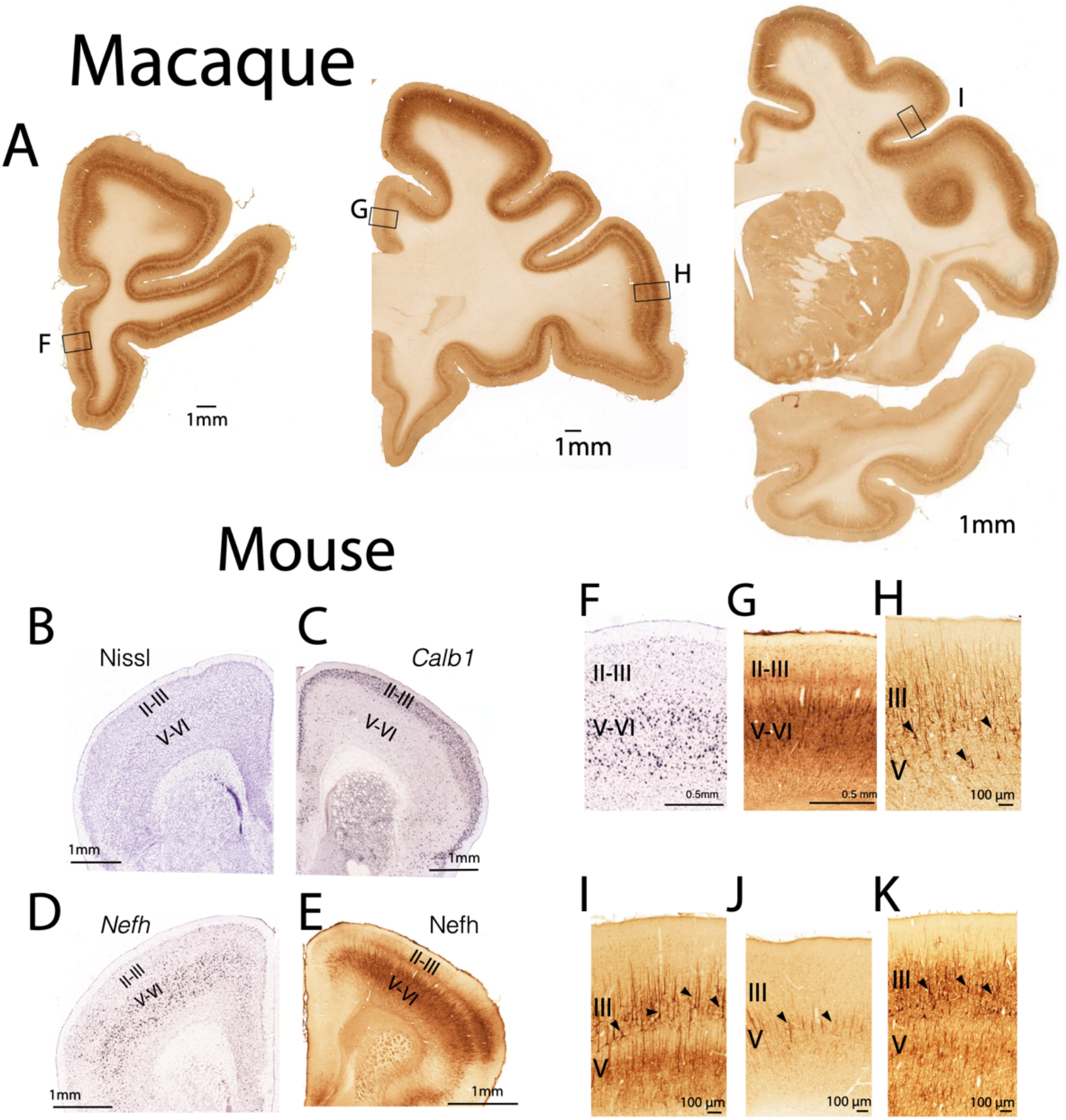
*NEFH* expression is greater in cortical layers II-III in the macaque compared with the mouse. The expression of *NEFH* is of particular interest because it is expressed by large neurons thought to project over long distances. Spatial variation in *NEFH* expression can be used to make inferences about modifications to connectivity patterns. Somas of large layers II-III neurons are expected to project cross-cortically whereas somas of neurons located in layers V-VI are expected to project subcortically. Accordingly, an increase in *NEFH* expression in layers II-III suggests an amplification of long-range cross-cortically projecting neurons. In macaques, (*A*) *NEFH* expression is relatively high in layers II-III and V-VI across the frontal cortex. This is in contrast to mice, which have relatively thin layers II-III. We used *Calb1* as a marker for layers II-III; the expression of *Calb1* spans layers II-III in mice. *NEFH* mRNA (*D*) and protein expression (*E*) is high in layers V-VI but extremely low in layers II-III in mice. (F–G) Close-up views through the mouse FC in *D* and *E* also show that *NEFH* mRNA (*F*) and protein (*G*) expression in layers II-III are particularly low in mice. This is in contrast to macaques, which show relatively stronger expression of NEFH protein in layers II-III across multiple frontal cortical areas (*H*–*K*) with the exception of the cingulate gyrus (*J*). These qualitative observations demonstrate major modifications in frontal cortex circuits between macaques and mice. The increased *NEFH* expression in layers II-III of macaques relative to mice suggests an expansion in cross-cortically projecting FC neurons may have emerged early in primate evolution.

## Supplementary tables

**Table S1**. Corresponding time points across humans and macaques.

**Table S2**. List of RNA sequencing datasets and age of samples used in the present study.

**Table S3**. Equations used to capture temporal trajectories in myelin water fraction (MWF)

**Table S4**. Cell clusters from the human primary motor cortex used to define layer II-III neurons and non-neuronal cells.

**Table S5**. List of LRP markers identified from associations between normalized gene expression and MWF.

**Table S6**. Post hoc Tukey tests between correlation coefficients of LRP markers across different regions and cortico-cortical pathway types across humans, macaques, and mice.

**Table S7**. Non-linear regressions used to compare growth of the PFC and FC white matter.

**Table S8**. Statistics on pathway types through the PFC and remaining FC in humans and macaques.

**Table S9**. Scanning parameters used for diffusion MR tractography of humans, macaques, and mice.

**Table S10**. Proportion of pathway types across the FC of humans, macaques, and mice. We only

**Table S11**. PFC measurements collected at various time points in macaques.

**Table S12**. Relative gene expression and thickness measurements across FC layers in humans, macaques, and mice.

**Table S13**. Statistics results from Tukey HSD tests to compare the relative thickness of FC layer II-III in humans, macaques, and mice.

**Table S14**. List of tract-tracer experiments used to compare diffusion MR tractography with tract-tracers.

## Notes

### Competing Interest Statement

The authors have declared no competing interest.

